# Tissue-specific landscape of protein aggregation and quality control in an aging vertebrate

**DOI:** 10.1101/2022.02.26.482120

**Authors:** Yiwen R. Chen, Itamar Harel, Param Priya Singh, Inbal Ziv, Eitan Moses, Uri Goshtchevsky, Ben E. Machado, Anne Brunet, Daniel F. Jarosz

## Abstract

Protein aggregation is a hallmark of age-related neurodegeneration. Yet, aggregation during normal aging and in tissues other than the brain is poorly understood. Here we leverage the African turquoise killifish to systematically profile protein aggregates in seven tissues of an aging vertebrate. Age-dependent aggregation is strikingly tissue-specific, and not simply driven by protein expression differences. Experimental interrogation, combined with machine learning, indicates that this specificity is linked to both protein-autonomous biophysical features and tissue-selective alterations in protein quality control. Co-aggregation of protein quality control machinery during aging may further reduce proteostasis capacity, exacerbating aggregate burden. A segmental progeria model with accelerated aging in specific tissues exhibits selectively increased aggregation in these same tissues. Intriguingly, many age-related protein aggregates arise in wild-type proteins that, when mutated, drive human diseases. Our data chart a comprehensive landscape of protein aggregation during aging and reveal strong, tissue-specific associations with dysfunction and disease.

**HIGHLIGHTS:**

- Tissue-specific protein aggregation is prevalent during vertebrate aging
- Both protein biophysical properties and tissue-specific protein homeostasis patterns impact aggregation
- A segmental progeria model with accelerated aging exhibits selectively increased protein aggregation in affected tissues
- Many aggregates that accumulate during physiological aging are linked to disease

## INTRODUCTION

Aging is accompanied by a decline in the control of protein synthesis, folding, conformational maintenance, and degradation, collectively known as protein homeostasis or ‘proteostasis’ (Hipp et al., 2019; Kaushik and Cuervo, 2015; Klaips et al., 2018; Lopez-Otin et al., 2013; Taylor and Dillin, 2011; Walther et al., 2015; Yang et al., 2019). Compromised proteostasis can lead to protein aggregation (Balch et al., 2008; Ben-Zvi et al., 2009; Bence et al., 2001; Huang et al., 2019). Indeed, protein aggregates are known to accumulate during normal aging in non-vertebrate species (yeast, worms, and flies), and this accumulation may at least in part be a result of proteostasis decline (Ciryam et al., 2013; David et al., 2010; Demontis and Perrimon, 2010; Huang et al., 2019; Walther et al., 2015). Protein aggregates have also been detected with age in the brain of killifish and mice (Kelmer Sacramento et al., 2020). However, a systematic, tissue-level understanding of protein aggregation in the context of natural aging is entirely absent in vertebrates. As a result, we still lack fundamental understanding of how protein aggregates form *in vivo*, whether they arise in tissues other than the brain, and whether protein aggregation is driven by protein autonomous kinetic and thermodynamic constraints, a decline in proteostasis, or some combination.

Most knowledge of age-related protein aggregation comes from studies of diseases of the nervous system. Proteostasis decline and the concomitant increase in protein aggregate load may explain why age is the primary risk factor for neurodegeneration (Braak et al., 2013; Kaufman et al., 2016; Lu et al., 2013; Prusiner, 2013). Although muscle weakness and cardiomyopathies have also been linked to protein misfolding in a few cases (Jiang et al., 2001; Vogler et al., 2018), the relationship between protein aggregation, aging, and disease remain poorly understood in most tissues.

To study vertebrate aging in a high throughput manner, we and others have developed the African killifish *Nothobranchius furzeri*, hereafter ’killifish,’ as a new model system (Cellerino et al., 2016; Genade et al., 2005; Harel et al., 2015; Hu and Brunet, 2018; Kim et al., 2016). The killifish is the shortest-lived vertebrate that can be bred in captivity, with a median lifespan of 4-6 months. It exhibits key age-dependent phenotypes and pathologies that are also observed in other vertebrates (including humans): muscle deterioration, fertility decline, cognitive decline, and neurodegeneration (Di Cicco et al., 2011; Harel et al., 2015; Matsui et al., 2019; Valenzano et al., 2006a; Valenzano et al., 2006b). Experimental interventions that extend mammalian lifespan, such as dietary restriction and pharmacological interventions, have a similar impact in the killifish (Terzibasi et al., 2009; Valenzano et al., 2006b). Importantly, genetic mutant killifish that model a human disease with segmental aging (telomerase deficiency) have been generated, and they exhibit premature deterioration in specific tissues (e.g., testis, gut) (Harel et al., 2015), similar to human patients. Thus, the killifish provides a unique platform to study aging and age-related diseases in vertebrates.

Here we harness the power of this emerging model of vertebrate aging in a systems-level investigation of protein aggregation during aging across seven different tissues. We find that age-dependent changes in protein aggregation are highly tissue-specific, driven not only by protein-autonomous biophysical properties but also by tissue-specific imbalances in proteostasis. Such proteostasis imbalance may be partially driven by co-aggregation of protein quality control machinery itself in a feed-forward loop during aging. Protein aggregation in telomerase mutant killifish is increased in the affected tissues that experience premature aging and involves components of the telomerase machinery. Intriguingly, many proteins that aggregate with age are linked to Mendelian diseases with degenerative phenotypes that have not previously been connected to protein misfolding. Collectively, our data chart a landscape of the demise of protein homeostasis in an aging vertebrate. They further define tissue-specific patterns of protein aggregation and deterioration of protein quality control networks that likely influence many age-related diseases beyond neurodegeneration.

## RESULTS

### The landscape of protein aggregation in seven tissues during vertebrate aging

The impact of physiological aging on protein aggregation remains poorly understood, especially in tissues other than the brain. Therefore, we systematically investigated protein aggregation in seven different tissues (brain, gut, heart, liver, muscle, skin, and testis) during normal aging in wild-type killifish and a telomerase-deficient disease model (Harel et al., 2015) (Figures 1A and 1B). To this end, we isolated protein aggregates in native conditions, adapting a differential centrifugation protocol (Chen et al., 2021; Kryndushkin et al., 2013; Kryndushkin et al., 2017) that physically separates high molecular weight protein aggregates (∼164 – 8804 Svedberg units, 1.03x10^6^ – 4.03x10^8^ kDa or 132-968 nm in diameter; STAR Methods) from the soluble proteome, large complexes, ribonucleoprotein particles, subcellular organelles, and large biomolecular condensates (e.g., stress granules, P-bodies; Figure S1A-D). To capture early aggregation events, including SDS-sensitive oligomers that have been linked to cytotoxicity (Chiti and Dobson, 2006; Hartl et al., 2011; Kayed et al., 2003), we avoided the use of ionic detergents (Figure S1A; STAR Methods). We validated this isolation procedure in *Saccharomyces cerevisiae*, where it effectively separated and identified aggregates of the proteins Sup35 and Rnq1 in cells harboring the prion forms of these proteins (Figure S1C; STAR Methods). It did not detect any aggregation of these proteins in genetically identical cells in which they were soluble (Figure S1C; Table S1).

**Figure 1.**
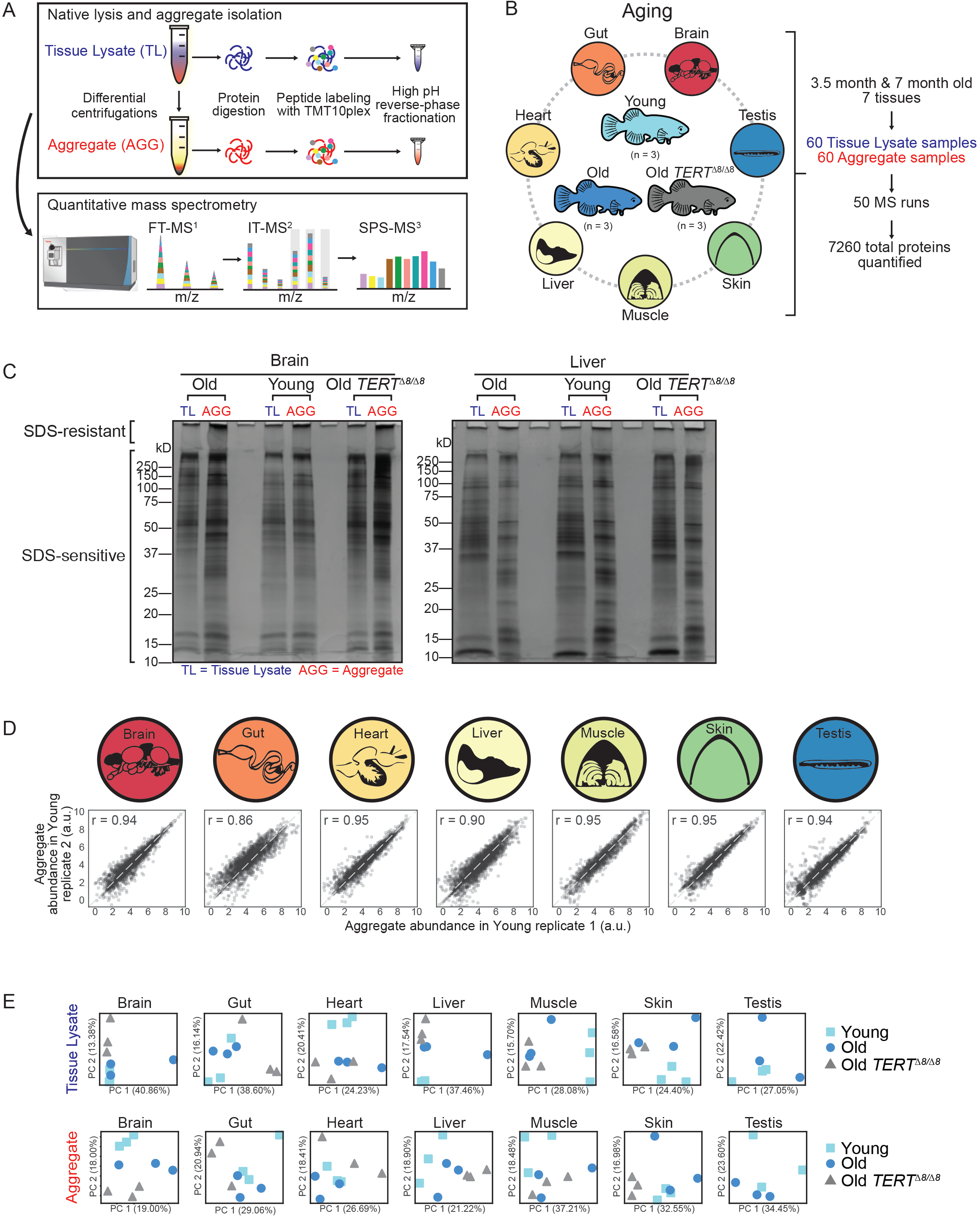
Quantitative proteomics of tissue lysate and aggregate fractions in young, old, and old telomerase mutant killifish. (A) Experimental workflow of tissue lysate extraction and isolation of high molecular weight protein aggregate fraction for tandem mass tag (TMT)-based quantitative mass spectrometry analysis. Killifish tissues were homogenized to isolate tissue lysates (TL) and a high molecular weight fraction enriched with protein aggregates (AGG). The TL or AGG fraction from each tissue was trypsin-digested, and the resulting peptides were labeled with 10plex-TMT isobaric tags, then subjected to high pH reverse phase fractionation, and analyzed on Thermo Orbitrap Fusion mass spectrometer. (B) Experimental design. Seven tissues (brain, gut, heart, liver, muscle, skin and testis) from 3 young (3.5 months), 3 old (7 months), and 3 old (7 months) *TERT^Δ8/Δ8^* male killifish were collected (except for testis in old *TERT^Δ8/Δ8^* killifish). Each tissue from an individual fish was homogenized to isolate the TL and AGG fractions (see Figure 1A). (C) Silver stain of tissue lysate and high molecular weight aggregate fractions from young, old, and old *TERT^Δ8/Δ8^* killifish brain and liver. Total tissue lysate (TL) and high molecular weight aggregate fraction (AGG) were resuspended in 5% SDS-sample buffer (without boiling), resolved by SDS-PAGE, and the gels were stained with silver stain. The brain image was also used in brain paper. (D) Reproducibility of aggregate abundance between biological replicates. Correlation between aggregate abundance (log2 transformed normalized peptide spectra counts) in AGG sample from Young fish #1 (x-axis) and Young fish #2 (y-axis) for each tissue, as quantified by TMT mass spectrometry analysis. Pearson’s correlation coefficient r is reported. Correlations between AGG samples from young fish #1 with young fish #3, young fish #2 with young #3, and similarly between 3 biological replicates for old fish and old *TERT^Δ8/Δ8^* fish are in Figure S1F. Similarly, correlations between TL samples from 3 biological replicates are in Figure S1F. Pearson’s correlation coefficients for all pairs of biological replicates are in Table S2B. Unit for y-axis a.u. stands for arbitrary units. (E) Principal component analysis (PCA) of protein abundance (log2 transformed normalized peptide spectra counts) in tissue lysate (TL) and high molecular weight aggregate fractions (AGG) in each tissue. Each symbol represents an individual fish: young (light blue squares), old (dark blue circles), old *TERT^Δ8/Δ8^* (grey triangles). The PCA for brain was also reported in the accompanying paper (Harel, Chen, et al.).

Using this isolation procedure in the killifish, we extracted both the tissue lysate (TL) and the aggregate fractions (AGG) from brain, gut, heart, liver, muscle, skin, and testis in three young (3.5 months) and three old (7 months) wild-type killifish, as well as from three old (7 months) telomerase-deficient (*TERT ^Δ8/Δ8^*) killifish (Figures 1A and 1B). Because telomerase-deficient killifish exhibit drastic age-dependent hypoplastic testis (Harel et al., 2015), we were unable to isolate protein from this tissue in *TERT^Δ8/Δ8^* animals. In total, we examined 120 samples (60 for tissue lysate and 60 for aggregate fraction) (Figures 1A and 1B), identifying increased protein aggregates with age in multiple killifish tissues, including the brain (Figure 1C). We next digested the proteins into peptides and labeled the 120 samples with isobaric Tandem Mass Tags (TMT10plex) to enable simultaneous quantification, treating each tissue and sample type (TL or AGG) as one set to minimize sample complexity. We performed high pH reversed-phase chromatography to reduce sample complexity further, and the resulting fractions were analyzed in 50 independent mass spectrometry runs (Figure 1B). Pooling samples in each labeling set as an additional TMT channel enabled robust identification and quantification of proteins across conditions (young, old, old *TERT^Δ8/Δ8^* individuals). Collectively, this suite of proteomics experiments detected 7,260 proteins across all 120 samples (6,169 in tissue lysates; 5,819 in aggregates) and on average ∼2,500 proteins for each sample using a stringent false discovery rate (FDR) of 1% (Figures 1B and Figure S1E; Table S2). We treated individual fish as separate samples to allow paired analysis across multiple tissues. The use of isobaric TMT tags and the pooled channel, together with a robust, optimized aggregate isolation protocol, enabled highly reproducible protein quantification between different animals for a given tissue (Pearson’s correlation coefficient r = 0.94+/-0.046 for the tissue lysate (TL) fraction, r = 0.92+/-0.028 for the aggregated (AGG) fraction; Figure 1D and Figure S1F; Table S2). Principal component analysis (PCA) readily separated samples according to age, disease, and tissue in both tissue lysate and aggregate fractions (Figure 1E). Thus, this systems-level proteomic experiment robustly identified and quantified both age- and tissue-dependent protein aggregation.

A fraction of proteins that aggregate in killifish overlapped with those identified in *C. elegans* (David et al., 2010; Walther et al., 2015), several of which also showed an age-associated increase (e.g., RHOA) (Figure S1G; Table S3-S4). Comprehensive studies of protein aggregation across aging tissues have not been performed to date in vertebrates, but one study did examine protein aggregation in aged mice brains and, at a lower coverage than in this study, aged killifish brains (Kelmer Sacramento et al., 2020). Our results overlapped well with both datasets (Figure S1G; Table S3). Thus, our approach identified previously known protein aggregates that arise during aging and, because of its extreme depth of coverage and examination of multiple tissues, also uncovered many new protein aggregates associated with natural vertebrate aging. Together, these quantitative profiles of protein aggregation and abundance in seven different tissues in aging wild-type and telomerase-deficient killifish provide a comprehensive landscape of protein aggregation during aging.

### Widespread tissue-specific aggregation in normal vertebrate aging

Proteins that exhibited increased aggregation with age were highly specific to each tissue (brain, gut, heart, liver, muscle, skin, testis), with only a few shared across tissues (Figure 2A, middle panel, and Figure 2B; Table S5). Similarly, proteins with increased aggregation propensity (amount of protein detected in the aggregate fraction normalized by total abundance in tissue lysate) were also tissue-specific (Figure 2A, right panel, and Figure 2B; Table S5). Consistent with previous mass spectrometry studies of the soluble proteome in aged mice and rats (Ori et al., 2015; Walther and Mann, 2011; Yu et al., 2020), most proteins were expressed similarly in young and old killifish across all tissues in our study (Table S2A). Thus, although some of the tissue-specific aggregation is expected based upon originates from tissue-specific expression profiles (Figures 2A, left panel, Figures S2A to S2B), the proteins that aggregate with age were far more specific to each tissue than changes in protein expression (Figure 2B-2C and S2C-S2D). Indeed, protein aggregation was not solely driven by changes in total protein amounts (Figure S2E): many proteins were expressed equivalently in multiple different tissues, yet only aggregated in one of them (Figure 2C). For example, during the course of aging, the long-chain fatty acid transporter SLC27A1 aggregated more in the brain than in the heart, the vacuolar sorting protein VPS35 aggregated more in the liver than in the heart, and the mitochondrial enzyme PPIF aggregated more in the skin than in the testis. Yet the proteins were similarly expressed in these tissues, and their levels did not change with age (Figure 2C). Similarly, Lamin A (LMNA) aggregation, and especially its aggregation propensity, increased more in the heart than the liver with age (Figure 2C, Figure S2F).

**Figure 2.**
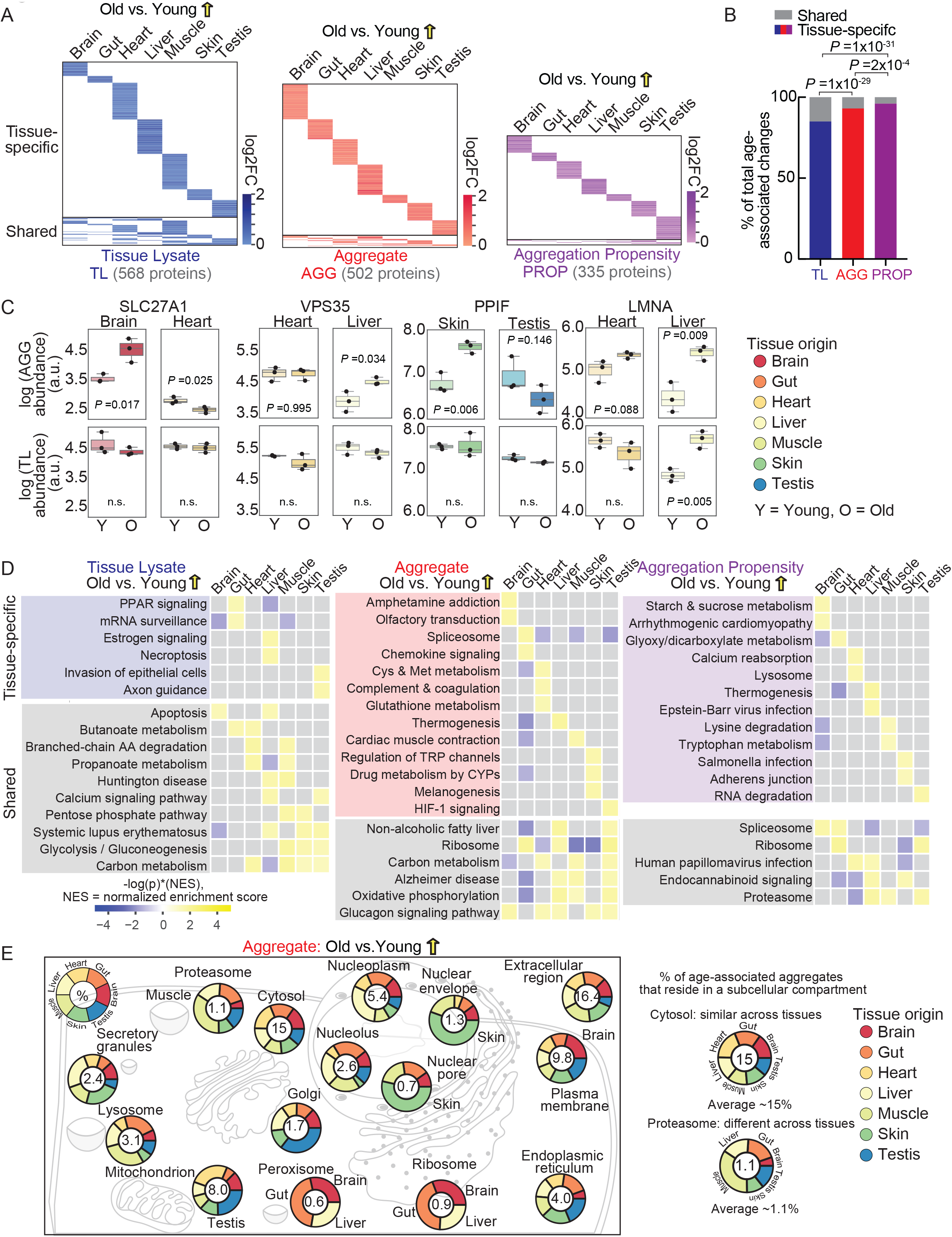
Tissue-specific changes in tissue lysate, aggregates, and aggregation propensity with age. (A) Heatmap on proteins that are significantly upregulated (yellow upward arrow, log2-transformed fold change in protein abundance (log2FC) > 0, two-sided Student’s t-test p-value < 0.05) with age (old over young) in tissue lysate (TL), aggregate (AGG), and aggregation propensity (PROP) across all tissues. Aggregation propensity is defined as the ratio of protein abundance in AGG over TL. The log2FC represents the average among 3 fish. Tissue-specific proteins (i.e. proteins with significant age-associated changes in only a single tissue) are at the top, and shared proteins (i.e. proteins with age-associated significant changes in at least two tissues) are at the bottom. The heatmaps are scaled based on total number of proteins with such significant positive differential changes (shown in bracket for TL/AGG/PROP). The identities of proteins significantly upregulated with age are listed in Table S5A-C. (B) Percentage of shared and tissue-specific changes among the entire dataset that showed a significant increase (log2-transformed protein abundance fold change (log2FC) > 0, two-sided Student’s t-test p-value < 0.05) in tissue lysate (TL) protein abundance, high molecular weight aggregate fraction (AGG) protein abundance, or aggregation propensity (PROP) in old compared to young fish. Shared and tissue-specific proteins were defined as in Figure 2A. P-values are from a Chi-squared test of independence on tissue-specific versus shared proteins across all tissues for TL and AGG, TL and PROP, and AGG and PROP. All data are available in Table S5D. (C) Examples of proteins with a tissue-specific increase in aggregate (AGG) in old fish compared to young fish despite similar tissue lysate (TL) abundance in different tissues. The box shows the quartiles of the data while the whiskers extend to show the rest of the distribution on protein abundance (y-axis) in young (Y) or old (O) samples. Each dot represents the protein abundance of an individual fish sample. Unit for y-axis a.u. stands for arbitrary units. P-values from Student’s t-tests are reported for all aggregates; n.s. indicates p>0.05. (D) Heatmap of significant functional and pathway enrichments identified among upregulated proteins in aging killifish using Gene Set Enrichment Analysis (GSEA). The proteins were ranked and sorted in descending order based on multiplication of log2-transformed fold change and –log10(p-value) between old versus young killifish (yellow upward arrow). Due to space constraints, only the top 3 (ranked by the highest normalized enrichment scores (NES)) significantly (p-value < 0.05) enriched Kyoto Encyclopedia of Genes and Genome (KEGG) terms in every tissue were shown for TL, AGG, and PROP. Tissues-specific terms were placed on top, whereas shared terms were placed at the bottom. The full lists of enrichment terms are available in Table S6. The color was scaled according to the rank statistic computed as the product of multiplying -log10(p-value) by NES. (E) Subcellular localization and complex association of proteins with significant accumulation of aggregate (AGG) in old tissues compared to young tissues (yellow upward arrow). Two examples (cytosol as a compartment with little tissue-specific differences and proteasome as a compartment exhibiting tissue-specific differences) were illustrated on the left. The arc lengths in the doughnut plot for each tissue are proportional to the respective fractions of aggregates that increase with age and reside in the compartment across tissues. The average value of such fraction across all tissues is reported in the center for each cellular compartment. Only tissues that contain proteins with a significant age-dependent increase in aggregates residing in the cellular compartments were visualized and counted towards the average calculation. On the right, doughnut plots are represented in proximity to the subcellular compartment of interest. Subcellular localization and complex association were inferred from the homologous human proteins retrieved from UniProt. Proteins that were annotated to exist in multiple compartments were double-counted. The fractions of upregulated aggregates residing in all compartments in each tissue are available in Table S7.

The tissue-specific patterns of protein aggregation impact many different biological functions (Figure 2D; Table S6). For example, the gene set enrichment analysis (GSEA) term linked to glutathione metabolism was enriched among proteins that aggregate more in the aging heart but not in other tissues (Figure 2D, middle panel; Table S6). Similarly, the GSEA term linked to thermogenesis was enriched among proteins that aggregate more in the aging liver but not significantly in other tissues (Figure 2D, middle panel; Table S6). The tissue selectivity of these enrichments was most apparent when considering aggregation propensity (Figure 2D, right panel; Table S6), further highlighting that tissue specificity of protein aggregation is largely independent of protein abundance. Moreover, these observations emphasize that although protein aggregation occurs during aging across many tissues, the alterations of specific biological functions likely differ substantially between them.

We next investigated the complex association and subcellular localization of proteins that significantly increased within the aggregate fraction of each aged tissue (Figure 2E and Figure S2G; Table S7). Analysis with the UniProt database for homologous human proteins revealed many tissue-specific patterns of subcellular localization associated with age-dependent protein aggregation (Figure 2E). Membrane proteins aggregated more commonly in the old brain than in any other tissue (Figure 2E). Nuclear pore and nuclear envelope proteins aggregated most prominently in the aging skin (Figure 2E). Specific ribosomal components aggregated in the old brain, gut, and liver (Figure 2E). Proteins associated with the proteasome and chaperonins commonly aggregated in the old gut, heart, liver, and testis (Figure 2E). Thus, although protein aggregation is a common feature in all aging vertebrate tissues, the precise identity of the aggregating proteins, and the subcellular compartments and biological pathways affected, differs substantially from one tissue to another.

### Biophysical features of age-associated protein aggregates

Two non-mutually exclusive possibilities could underlie the tissue specificity of aggregates: i) the biophysical properties of specific proteins expressed in a given tissue could differ (and, hence, the inherent propensity of these proteins to aggregate, which may put them more at-risk during aging), ii) the protein quality control network could deteriorate differently in different tissues during aging. To test the first possibility, we analyzed 37 protein features (STAR Methods), including many that have previously been associated with protein aggregation, such as intrinsic protein disorder, charge distribution, aromatic residue enrichment, and others across the entire killifish proteome (Figure 3A; Table S8). We verified that no specific enrichment biases in biophysical features were present in any tissue, even when accounting for relative protein abundance (Figure S3A-S3B; Table S8). Indeed, in young animals the biophysical features enriched in aggregation-prone proteins were largely shared among different tissues (Figure 3B; Table S8).

**Figure 3.**
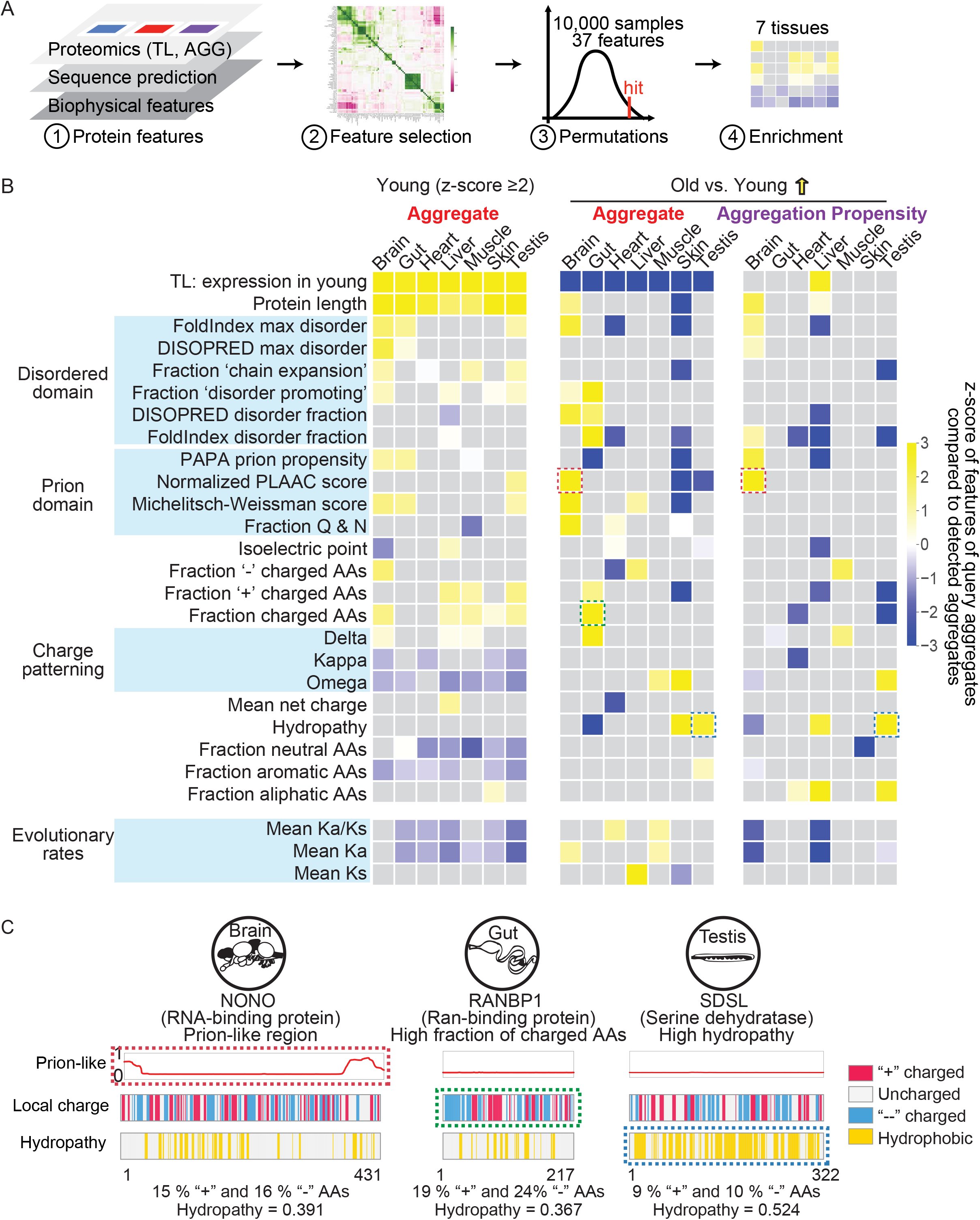
Features of proteins that are aggregation-prone in an age-dependent manner across tissues. (A) A schematic representation of the analysis workflow to characterize features of the proteins detected in tissue lysate (TL) and aggregate fractions (AGG) as well as those with age-associated changes. We included protein abundance data from this proteomic study as well as features calculated using various computational tools. We next conducted pair-wise correlation analysis on these features, eliminated a subset of features that were highly correlated with others, and focused on 37 key features, including many previously associated with protein aggregation, such as intrinsic protein disorder, charge distribution, aromatic residue enrichment, and others across the entire killifish proteome. We conducted Monte Carlo simulations (sample 10,000 times with replacement) to identify sequence feature enrichment and assign statistical significance. (B) Biophysical features of proteins that contributed the most (expression z-score ≥2) to aggregate burden in young animals and proteins with significant (Student t-test p-value < 0.05) age-associated increase (yellow upward arrow) in aggregate (AGG) and aggregation propensity (PROP). The features we investigated include “FoldIndex max disorder” – the maximum stretch of disorder predicted by FoldIndex (Prilusky et al., 2005), “DISOPRED max disorder” – the maximum stretch of disorder predicted by DISOPRED (Jones and Cozzetto, 2015), “Fraction chain expansion” – the fraction of residues that contribute to chain expansion (E/D/R/K/P) (Holehouse et al., 2017), “Fraction disorder promoting” – fraction of residues that is predicted to be ‘disorder promoting’ in TOP-IDP-scale (Campen et al., 2008), “DISOPRED disorder fraction” – the fraction of total residues with a DISOPRED score above 0.5, “FoldIndex disorder fraction” – the fraction of total disorder residues predicted by FoldIndex, “PAPA prion propensity” – the predicted prion propensity by PAPA (Toombs et al., 2012), “Normalized PLAAC score” –the normalized prion score NLLR predicted by PLAAC (Lancaster et al., 2014), “Michelitsch-Weissman score” – prion score predicted by methods developed by the Weissman lab (Michelitsch and Weissman, 2000), fraction of Q and N residues, protein isoelectric point pI, fraction of negatively charged residues, fraction of positively charged residues, fraction of charged residues, protein charge patterning parameters ‘delta’, ‘kappa’, and ‘Omega’ (Das and Pappu, 2013), mean net charge, protein hydropathy, fraction of neutral, aromatic, and aliphatic residues, non-synonymous evolutionary rates Ka, synonymous evolutionary rates Ks, and their ratios Ka/Ks. A one-tailed statistical test was performed for each tissue where aggregates from the tissue were sampled 10,000 times (with replacement), drawing exactly the same number of proteins as the query set from a given tissue. Terms were visualized only when the average feature value from the query set of proteins were significantly (one-tail test at 0.05 cutoff) different from the Monte Carlo simulation derived population distribution, whereas the nonsignificant ones were in gray. The color in the heatmap reflected the z-score of the average feature value from the query set compared to the Monte Carlo simulation-derived test population means. See STAR Methods for details on the implementation and analysis of the Monte Carlo simulation. A few tissue-specific features of proteins with increased aggregate or aggregation propensity are highlighted with the dotted box in the heatmap. Example protein sequence feature profiles are available in Figure 3C. See Table S8A-D for results. (C) Biophysical properties of example proteins with significantly increased aggregate (AGG) or aggregation propensity (PROP) with age. The prion-like region (predicted by PLAAC (Lancaster et al., 2014)), net charge per residue (localCIDER with a sliding window of 5 residues (Das and Pappu, 2013)), and hydropathy based on Kyte-Doolite scale (localCIDER with a sliding window of 5 residues) are illustrated for NONO, RANBP1, and SDSL. Residues predicted to have a positive net local charge (NCPR>0) are shown as red bars, whereas negative net local charges are shown as blue bars (middle). Residues with local hydropathy scores larger than 0.5 are shown as yellow bars (bottom). Tissue-specific features (Figure 3B) are highlighted with a dotted box. The fraction of residues that are positively charged, negatively charged, and hydrophobic are listed underneath each protein.

In contrast, no single feature predicted either age-associated changes in aggregate or aggregation propensity across tissues (Figure S3C). Instead, age-dependent aggregates were enriched in very different biophysical features depending upon the tissues in which they arose (Figure 3B). For example, proteins harboring long intrinsically disordered regions and prion-like sequences were significantly more likely to aggregate with age in old brains (Figure 3B-3C, e.g., NONO, see also other examples in the accompanying paper). However, enrichment of charged residues strongly correlated with age-dependent increases in aggregation in the gut (Figure 3C, e.g., RANBP1). Likewise, higher hydropathy (Kyte and Doolittle, 1982) was a strong predictor for age-dependent aggregation and aggregation propensity in the testis (Figure 3C, e.g., SDSL). Importantly, these biophysical enrichments did not simply reflect biases in the expressed proteome of any tissue that arose during aging (Figure S3A-B). Instead, our data establish that distinct biophysical predictors of protein aggregation emerge within a tissue during aging. That is, even the protein-autonomous signatures of age-dependent aggregation are highly tissue-specific.

### Contributors to protein aggregation *in vivo*

The protein-autonomous signatures that we observed led us to suspect that many proteins we identified have an intrinsic ability to aggregate when expressed. To test this prediction, we turned to *Saccharomyces cerevisiae*, a widely used model for protein quality control and aggregation (Figure 4A). We selected 47 proteins that aggregated with age in different killifish tissues, tagged them with eGFP, expressed them in yeast, and measured the number of cells with fluorescent puncta (an indication of protein aggregation; Figures 4A and S4A). In these experiments 27 of the 47 killifish proteins (57%) aggregated (Figures 4B and 4C; Table S9), including NCOA5 (brain), MRPS18B (gut), ADCK3 (heart), ECHS1 (liver), DHTKD1 (muscle), CALM3 (skin) and PDHB (testis). By contrast, eGFP did not aggregate under these conditions. Because of slight differences in laboratory husbandry temperatures for killifish (26 °C) and *S. cerevisiae* (30 °C), we also repeated some of these experiments in yeast at 26 °C and observed no difference in aggregation behavior (Figure S4B). Likewise, aggregation did not correlate with fluorescence intensity (Figure S4C), as would be expected if these differences were an artifact of differences in protein levels. Thus, many, but not all, proteins that aggregate with age in killifish appear also to have an intrinsic capacity to do so even when expressed in the context of a youthful yeast proteome.

**Figure 4.**
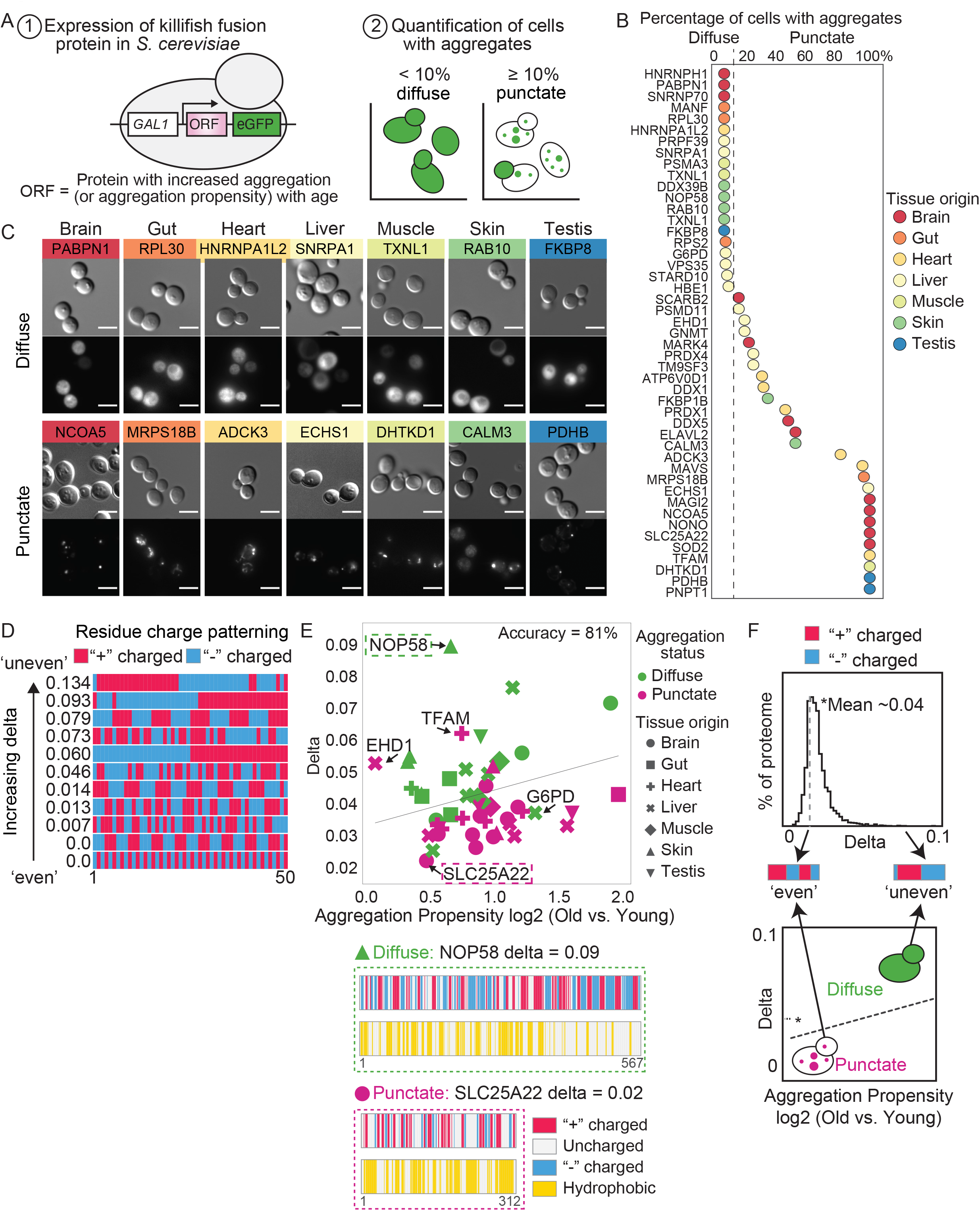
Validation of protein aggregation and identification of contributors to protein aggregation *in vivo*. (A) Schematic representation of the *S. cerevisiae* aggregation assay (1) and criteria used to score intrinsic aggregation behavior based on fractions of cells with aggregated eGFP morphology (2). Killifish proteins of interest (open reading frame, ORF) fused with eGFP were expressed exogenously in yeast under a GAL1 promoter that is inducible upon switching the sugar source of the yeast growth media from raffinose to galactose. The percentage of cells with aggregates (fluorescent puncta) was counted. A protein is considered diffuse if less than or equal to 10% of the cells expressing the eGFP-fusion protein contains puncta. A protein is considered to be in punctate form if more than 10% of the cells expressing the eGFP-fusion protein contains puncta. (B) Quantification of the percentage of cells with aggregates assessed upon overexpression in *S. cerevisiae* as described in Figure 4A. Proteins with a significant age-associated increase in aggregate and/or aggregation propensity (Student’s t-test p-value <0.05, log2(fold change) in AGG or PROP >0) in old versus young samples were selected, and their tissue origins (all tissue-specific) are differentiated by the colors of the dots. Proteins were ranked based on the fraction of cells with puncta from 0 to 100%. The dashed line represented the 10% cutoff that was used to call diffuse (<= 10%) versus punctate (>10%) proteins. Two independent *in vivo* aggregation experiments were performed at 30°C and an average of ∼120 EGFP-positive cells were quantified for each protein in a given experiment. The quantification results are available in Table S9. (C) Example microscopy images of the proteins of interest (with a significant age-associated increase in aggregate and/or aggregation propensity with age) upon overexpression in *S. cerevisiae* as outlined in A and quantified in B. Two proteins, one diffuse and one punctate, based on the *in vivo* assay, were chosen from each tissue. The images chosen were representative of two independent experiments. (D) Examples of peptide sequences with varying degrees of charged residue mixing and the corresponding delta scores (computed according to (Das and Pappu, 2013)). (E) A support vector machine classifier on *in vivo* aggregation status of age-associated killifish aggregates (increased aggregation propensity in old over young fish) assessed by yeast overexpression assay outlined in Figure 4A. Age-associated aggregation propensity change and charge asymmetry deviation within a sequence (delta computed in localCIDER) for each protein are shown in x and y axis respectively and their *in vivo* aggregation status can be distinguished by color. Tissues where the protein exhibited increased aggregation propensity changes are distinguished by the different marker shapes. The solid line separates those that showed diffuse eGFP morphology upon protein overexpression. Sequence feature profiles of two proteins (one classified as diffuse whereas the other as punctate) were generated as in Figure 3C. See STAR Methods for details. (F) The distribution of delta values from all aggregates detected in our study and a model that predicts proteins aggregation behavior *in vivo* based on charge patterning ‘delta’ and changes in protein aggregation propensity during aging.

Next, we endeavored to determine features distinguishing proteins that aggregated in these experiments from those that did not. We built a support vector machine classifier model for the aggregation status for each protein in yeast, considering two features at a time to avoid overfitting (STAR Methods). Previous classifier models that are standards in the field separate “globular folded” and “natively unfolded” proteins and have been built mainly from *in vitro* data (Uversky et al., 2000). These were unable to discriminate the proteins that aggregated in our assay from those that did not (Figure S4D). By contrast, our best classifier was able to discriminate between these classes of proteins based on two features: a charge patterning parameter (‘delta,’ which measures the amount of charge and charge asymmetry across a sequence (Figure 4D) (Das and Pappu, 2013; Holehouse et al., 2017) and age-related changes in aggregation propensity in killifish (Figure 4E; Table S10).

Interestingly, killifish proteins that did not aggregate when expressed in yeast had two defining characteristics. First, they had unusually skewed patterns of charge distribution within the polypeptide sequence. Second, they were generally devoid of long hydrophobic stretches (Figure 4E-F). Our second-best model classified killifish proteins that aggregate in the yeast system based on the maximum number of asparagine (Q) and glutamine (N) residues in a sliding window and the ‘delta’ charge patterning parameter (Figure S4E-G). This is intriguing because both features were enriched in aggregation-prone proteins from young killifish (Figure 3B) and point to their important role as intrinsic determinants of protein aggregation behavior. Together, our findings suggest that protein-autonomous properties, especially the local electrostatic and hydrophobic environment, influence, at least in part, aggregation status in the context of aging.

### Tissue-specific changes in the proteostasis network

Could changes in the proteostasis network (e.g., chaperones, proteasome) contribute to tissue-specific aggregation with age? This idea has often been invoked, but our dataset provides a unique opportunity to investigate this question from the standpoint of chaperones and clients alike (Higuchi-Sanabria et al., 2018; Hipp et al., 2019; Kaushik and Cuervo, 2015; Morimoto, 2020; Pilla et al., 2017; Pras and Nollen, 2021; Taylor and Dillin, 2011). Although we observed relatively few age-dependent changes in the total lysate proteome, many of those we did see occurred in components of the proteostasis network (Figures 5A and S5A-B; Table S11). For example, in old animals, proteasomal proteins were downregulated in the liver (Figure 5A and Figure S5B; Table S11), and several chaperone proteins were downregulated in both liver and muscle (Figure 5A and Figure S5A; Table S11). Despite the lack of comprehensive proteomic datasets investigating vertebrate aging, comparisons to studies of individual tissues in other animals indicated a good degree of overlap. For example, in old killifish levels of the Hsc70/HSPA8 chaperone decreased in brain, skin, testis, and muscle (Figure S5A), but increased in the heart (Figure S5A). Similar changes in HSPA8 expression have been observed in targeted studies in mouse (brain and heart) (Walther and Mann, 2011) and human (muscle) (Murgia et al., 2017) tissues. In addition, the heart-specific small heat shock protein HSPB7, which is essential for maintaining myofibrillar integrity (Golenhofen et al., 2004), strikingly decreased with age in old killifish heart (Figure S5A). Finally, the levels of the myosin chaperone UNC45B (Price et al., 2002) and actin chaperone TRiC decreased drastically in aging killifish muscle (Figure S5A; Table S11).

**Figure 5.**
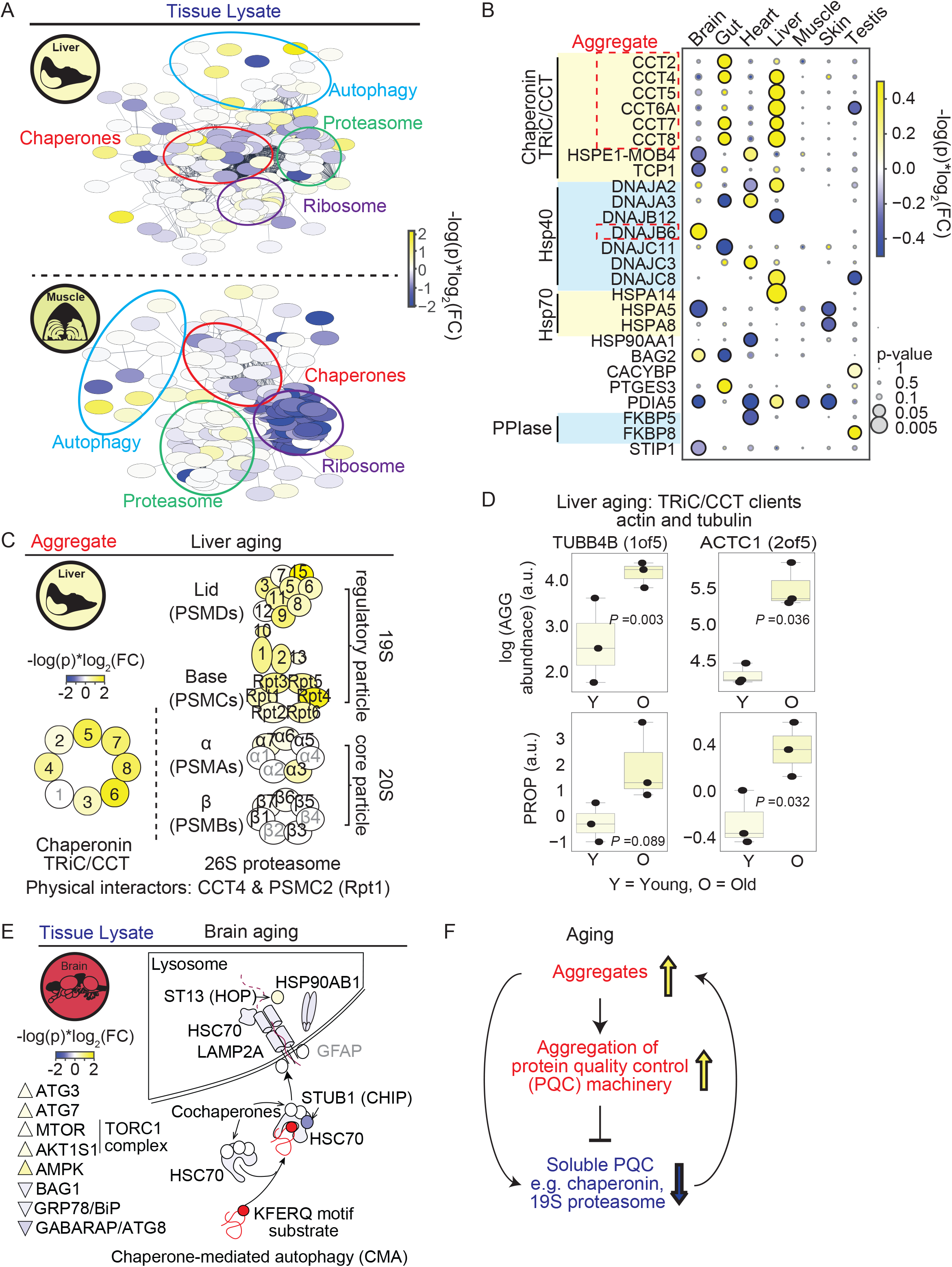
Tissue-specific changes in protein quality control machinery in tissue lysate and aggregate fractions during aging. (A) Protein quality control machinery shown as protein-protein interaction networks and their tissue lysate changes during aging in brain, liver and muscle. Proteins that are part of protein quality control machinery (ribosome, proteasome, and chaperones) or involved in ubiquitin proteolytic pathway (hsa04140 from KEGG database) and autophagy/lysosomal degradation pathway (hhsa04120 from KEGG database) were selectively analyzed. The interaction network of these proteins was inferred from the interactions of human homologs retrieved from STRING database (Szklarczyk et al., 2019) and visualized in Cytoscape software (high confidence setting was used where the minimum required interaction score was 0.7 out of a normal scale of 0 to 1). The color for each protein reflects the rank-statistics of its age-dependent tissue lysate change (defined by the multiplication of negative log10-transformed Student’s t-test derived p-value and log2-transformed fold change in tissue lysate in old samples compared to the young samples). The colored circles denote clusters of proteins from the same protein complex or involved in specific protein degradation pathways. (B) Tissue lysate abundance changes of proteins involved in chaperone-mediated autophagy (CMA) pathway and the changes of CMA-selective clients versus non-client in aging brain. The triangle/ellipse color for each protein is indicative of its ranked statistics (-log10-transformed p-value multiply by log2-transformed fold change of old over young). Proteins in light gray label were not detected in the tissue lysate fraction in brain. CMA-clients are proteins detected in killifish brain with peptide motif that meet the following criteria (Kaushik and Cuervo, 2018): include a glutamine on one of the sides and contains one or two of the positive residues K and R, one or two of the hydrophobic residues F, L, I or V and one of the negatively charged E or D residues. The log2-transformed fold change in tissue lysate and aggregate abundance of clients and non-clients in old versus young killifish brain were separately plotted (each dot represents a single detected protein). A boxplot is overlayed for each category and shows the quartiles of the dataset while the whisker extends to show the rest of the distribution. Independent two-sided t-tests were performed between clients and non-client for each sample type and the p-values were reported. (C) Chaperone abundance changes in high molecular weight aggregate fraction between old and young animals across all tissues. The circle color is indicative of the ranked statistics (-log10-transformed p-value multiply by log2-transformed fold change of old over young protein abundance) and the circle size is indicative of the -log10-transformed Student’s t-test p-value. Results are available in Table S11. (D) Aggregate abundance changes of TRiC and proteasome components and their clients in the liver with age. The circle/ellipse color for each protein is indicative of its ranked statistics (-log10-transformed p-value multiplied by log2-transformed fold change of old over young). Proteins in light gray labels were not detected in the aggregate fraction in the liver. (E) Aggregate and aggregation propensity changes of TRiC clients actin and tubulin during liver aging. The box shows the quartiles of the data while the whiskers extend to show the rest of the distribution on protein abundance (y-axis) in young (Y) or old (O) samples. Each dot represents the protein abundance of an individual fish sample. P-values are from Student’s t-test. (F) Model of a vicious cycle that drives protein aggregation during aging. Unfolded and misfolded protein aggregates sequester protein quality control (PQC) machinery such as chaperones and proteasome subunits into aggregate and effectively titrate away total PQC components. Decreased availability of PQC compromises cellular proteostatic capacity, which contributes to a higher aggregate burden and a further decline in proteostasis.

Several chaperones with significant changes in the aging brain are involved in chaperone-mediated autophagy (CMA). CMA is a selective degradation process in which proteins with a KFERQ-like motif are delivered to the lysosomes via HSC70 and co-chaperones and then internalized in lysosomes by the receptor lysosome-associated membrane protein type 2A (LAMP2A) (Kaushik and Cuervo, 2018). In old killifish brains, the CMA co-chaperones (e.g., carboxyl terminus of HSC70-interacting protein (CHIP), STUB1) were downregulated, whereas the HSP70–HSP90 organizing protein (HOP), ST13 was upregulated (Figure 5B and Figure S5A). In addition, total HSC70 (HSPA8) and HSP90 (HSP90AB1) levels were reduced along with LAMP2A (Figure 5B and Figure S5A), suggesting that the aging brain has reduced capacity for CMA. Consistent with this hypothesis, we observed a significant accumulation of CMA-substrates in the aggregate fraction from old brains that was not simply driven by changes in their expression level (p=3.2x10^-7^; Figure S5C). The increased aggregate burden of CMA substrates is specific to the aging brain (Figure S5C) and exemplifies how tissue-specific proteostasis collapse can cause tissue-specific protein aggregation with age.

Chaperones can co-aggregate with misfolded client proteins (Auluck et al., 2002; Cox et al., 2018; Fonte et al., 2002). We investigated whether this might contribute to age-dependent defects in proteostasis in killifish. We noted many chaperones in the aggregate fraction, and their presence was highly tissue-specific (Figure 5C and Figure S5D). Intriguingly, in many cases, the chaperone selectivity for its clients was connected to the biophysical predictors of aggregation for a given tissue. For example, we observed age-dependent increases in the aggregation of DNAJB6, an Hsp40 chaperone that acts on Q/N-rich substrates (Hageman et al., 2011), in the old brain (Figure 5C), and Q/N-rich prion-like sequences were highly enriched in age-induced protein aggregates from this same tissue (see Figure 3C). Likewise, similarities in the composition of chaperone proteins in the aggregate fraction between tissues can predict similarities in the aggregated proteomes of these tissues. For example, TRiC/CCT complex subunits accumulated in the aggregate fraction of both aged gut and liver (Figure 5C), and the types and abundances of proteins in the aggregate fraction from the aging gut and liver correlated more closely together than with any other tissues (Figure S5D). Moreover, additional protein quality control factors (e.g., peptide disulfide oxidoreductases PDIA4, PDIA6, and P4HB) were also among the strongest drivers of tissue separation in PCA analysis of age-dependent aggregates (largest PC2 loadings in Figure S5D; Table S12). Finally, titration of proteasome components may also contribute to differences in protein aggregation during aging. We found that the 19S regulatory particle of the proteasome was reduced in old liver tissue lysate (Figure S5B; Table S11) whereas it increased in aggregates from the same tissue (Figure 5D; Table S11). This may be a consequence of reduced TRiC/CCT levels, which would be expected to drive accumulation of unfolded nascent client polypeptides such actin and tubulin. Indeed, these proteins also accumulated in the aggregate fraction in aged animals (Figure 5E). Our data suggest that, in addition to protein biophysical properties, tissue-specific proteostasis breakdown is a key driver of protein aggregation with age. This breakdown may often involve aggregation of protein quality control factors themselves, sparking a ‘vicious cycle’ that further reduces proteostatic capacity.

### Protein aggregation in a model for a segmental age-dependent disease

Progeria often manifests segmentally, with features of premature aging emerging in specific tissues (Ullrich and Gordon, 2015). For example, dyskeratosis congenita – a disease that arises from telomerase deficiency – is characterized by premature aging-like phenotypes in highly proliferative tissues (e.g., gut, blood, skin, and testis) (Kirwan and Dokal, 2009). Like human patients, killifish *TERT^Δ8/Δ8^* mutants also experience a premature collapse of these tissues (Harel et al., 2015). Telomerase deficiency is known to be associated with genome instability (O’Sullivan and Karlseder, 2010), but whether it could also be accompanied by protein aggregation defects is unknown. To address this question, we analyzed aggregates in old *TERT^Δ8/Δ8^* animals that were age-matched with the wild-type fish we previously characterized. Proteins that aggregated in old *TERT^Δ8/Δ8^* animals were also highly tissue-specific both in protein identities and in the biological pathways in which they participate (Figure 6A-6B and Figure S6A; Table S14). The biophysical features correlated with protein aggregation in these animals were also diverse and depended on the tissue (Figure S6B). For example, proteins that aggregated in the liver of old telomerase mutants were enriched in prion-like domains relative to wild-type. Another prominent feature is an enrichment of intrinsically disordered regions in aggregates in the skin of old telomerase mutants (Figure S6B; Table S15). In addition, the fraction of positive charges and skewed charge pattern (delta) were enriched in protein aggregates in the skin of old *TERT^Δ8/Δ8^* individuals (Figure S6B; Table S14).

**Figure 6.**
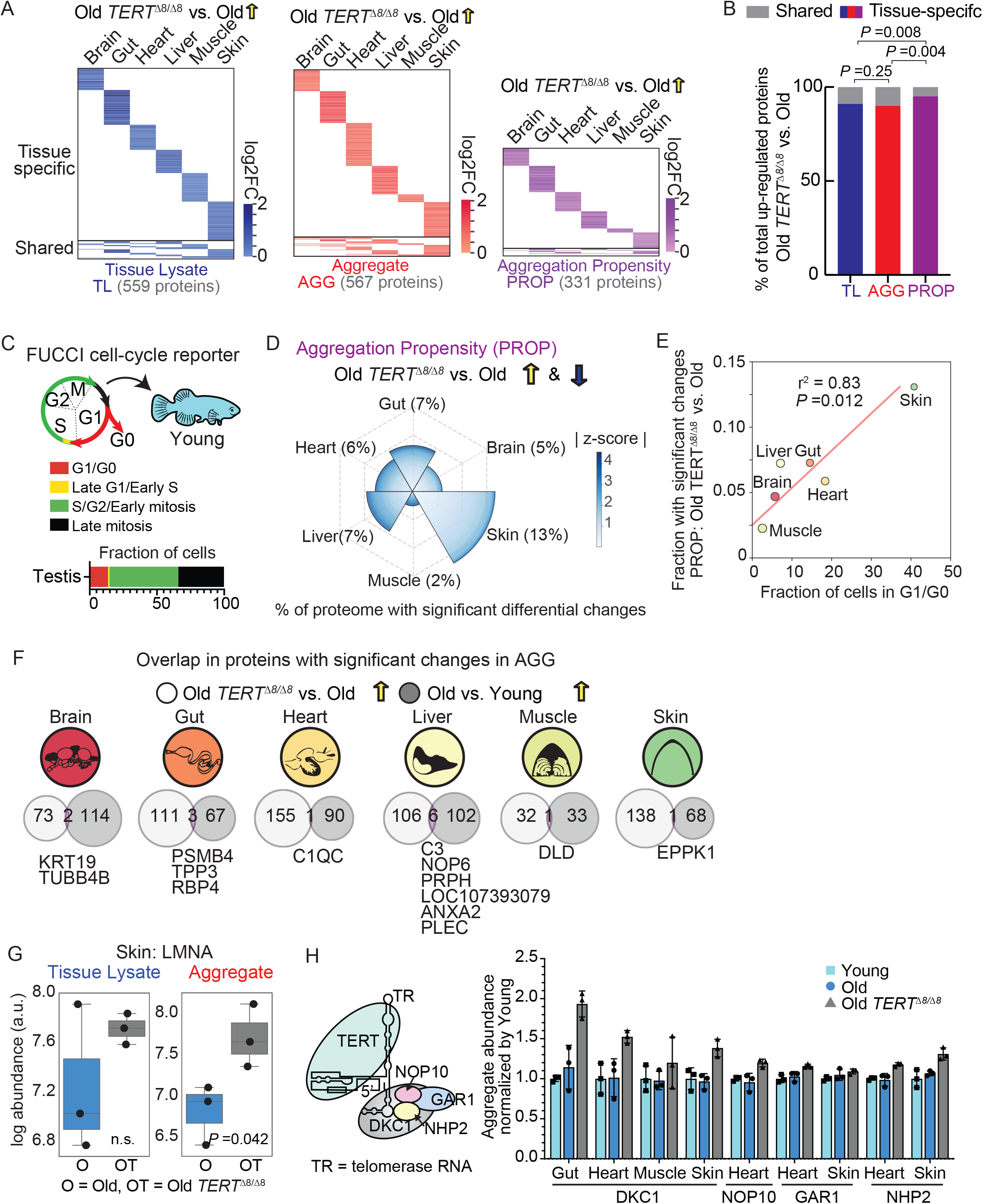
Analysis of changes in tissue lysate and high molecular weight aggregate fraction of old wild-type and old *TERT*^Δ8/Δ8^ mutant fish. (A) Heatmap on proteins that are significantly (Student’s t-test p-value < 0.05) upregulated in tissue lysate (TL), aggregate (AGG), and aggregation propensity (PROP) across all tissues in Old telomerase deficient (*TERT^Δ8/Δ8^*) killifish compared to its age-matched wild-type control fish. Tissue-specific (proteins with significant telomerase deficient-associated changes in only a single tissue) changes are placed at the top and shared (proteins with significant changes in at least two tissues) are placed at the bottom. The log2FC represented the average among 3 fish. The heatmaps were scaled based on the total number of proteins with significant positive differential changes (shown in brackets for TL/AGG/PROP). The identities of proteins significantly upregulated with age are in Table S13. (B) Percentage of shared and tissue-specific changes among the entire dataset that showed a significant increase (log2-transformed protein abundance fold change log2FC > 0, two-sided Student’s t-test p-value < 0.05) in tissue lysate (TL) protein abundance, high molecular weight aggregate fraction (AGG) protein abundance, or aggregation propensity (PROP) in Old *TERT^Δ8/Δ8^* mutants compared to age-matched wild-type fish. Shared and tissue-specific proteins were defined as in Figure 6A. P-values are from a Chi-square test of independence on tissue-specific versus shared proteins across all tissues for TL and AGG, TL and PROP, and AGG and PROP were reported. Data are from Table S13D. (C) Cell cycle progression inferred from adult male transgenic killifish carrying the cell-cycle dual FUCCI reporter (Dolfi et al., 2019). Three 2.5 months old fish were dissected, and the bulk population of each tissue underwent FACS sorting to infer the relative distribution of different cell cycle stages. The average results among the 3 biological replicates for testis were shown as an example. Results are available in Table S16. (D) Percentage of proteins with differential changes (include both up-regulation and down-regulation) in aggregation propensity (PROP) between age-matched old *TERT*^Δ8/Δ8^ and wild-type animals. The z-scores of changes in PROP between age-matched old *TERT^Δ8/Δ8^* and wild-type were calculated for each protein (i.e., TvO_prop_logFC was the log2-transformed fold change of old *TERT^Δ8/Δ8^* divided by age-matched wild-type) at each tissue level. Those that were significant (p-value <0.05) were counted, and the percentage of the proteome they represent is reflected in the radius of the sector for each tissue. The heatmap color is reflective of the ranked z-score of these significantly differentially regulated proteins with a higher degree of remodeling denoted with darker shades of blue. (E) Correlation between the fraction of cells in G1/G0 phase from bulk tissue population and the extent of aggregate remodeling across tissues during segmental aging due to telomerase deficiency. The fraction of cells in G1/G0 phase (x-axis) from each tissue is estimated based on the FUCCI reporter described in Figure 6C. The extend of aggregate remodeling (y-axis) is measured as the fraction of proteins detected in the aggregate fraction with significant differential changes (include both up-regulation and down-regulation) in aggregation propensity (PROP) between old *TERT^Δ8/Δ8^* vs. old wild-type (WT). Results are available in Table S16. (F) Overlap among aggregates that are significantly up-regulated in old compared to young (p-value <0.05, log2 (fold change) >0) and old *TERT^Δ8/Δ8^* mutants compared to age-matched wild-type (p-value <0.05, log2 (fold change) >0). The overlapped proteins were listed for each tissue. (G) Box plot of Lamin A protein level in skin samples obtained from Old *TERT^Δ8/Δ8^* mutant and age-matched wild-type fish. The box shows the quartiles of the data while the whiskers extend to show the rest of the distribution on protein abundance (y-axis) in old (O) or old *TERT^Δ8/Δ8^* (T) samples. Each dot represents the AGG abundance from an individual fish. P-values are from Student’s t-test; n.s. indicates p>0.05. (H) Illustration of components of the telomerase complex (left) and bar plot representation of normalized dyskerin (DKC1) level in the aggregate fraction in age-matched old *TERT*^Δ8/Δ8^ and wild-type killifish (right). The height of the bar plot represents the average, and the error bar represents the standard deviation. Results are available in Table S17.

We next compared tissue-specific aggregation patterns in the old *TERT^Δ8/Δ8^* to those of age-matched wild-type animals. If protein aggregation is central to vertebrate aging, we should see enhanced aggregation in the proliferative tissues selectively affected by segmental aging in this model. To quantify proliferation in multiple tissues, we used a transgenic killifish expressing a Cdt1-RFP-Geminin-GFP FUCCI cell cycle reporter (Dolfi et al., 2019) (Figure 6C and Figure S6C). With the exception of the heart, which is known to be more proliferative in fish than in mammals (Jopling et al., 2010; Wang et al., 2020), the proliferation capacity of the tissues we examined corresponded with intuition from prior literature (Figure S6C-S6D). Although the largest individual changes in aggregation occurred in the brain, few proteins were affected in this non-proliferative tissue (4-7%, Figure 6D; Table S2A; Table S16). By contrast, proliferative tissues such as skin and gut experienced far more protein aggregation events—with a larger fraction of proteome conferring significant changes in aggregation and aggregation propensity— in old *TERT ^Δ8/Δ8^* mutant compared to old wild-type killifish (12-17%, Figure 6D-E; Table S16). Taken together, our data emphasize that proliferative tissues of TERT*^Δ8/Δ8^* animals experience a greater extent of aggregate remodeling, with a larger fraction of the proteome being affected.

The increased number of aggregation events in TERT*^Δ8/Δ8^* mutant animals did not generally come from further accrual of protein aggregates that were already detected in old animals (Figure 6F), aside from several counterexamples such as complement component 3 (C3) and keratin 19 (KRT19) (Figure S6F). A previous study that modeled cellular senescence reported up-regulation of Progerin, a truncated form of LMNA that forms detergent-insoluble aggregates (Cao et al., 2011b), in human fibroblasts undergoing progressive telomere damage (Cao et al., 2011a). Notably, in old *TERT ^Δ8/Δ8^* killifish skin, LMNA aggregation was up by 85% compared to old wild-type skin with no significant increase in tissue lysate expression (Figure 6G).

Finally, some of these aggregated proteins were logically connected to the TERT mutant itself: proteins that interact with TERT and are involved in telomere repeat extension. For example, in aged *TERT^Δ8/Δ8^* mutants, the telomerase accessory component dyskerin protein (DKC1) showed significant elevation in the aggregate fraction of every tissue except muscle (not detected in brain, Figure 6H; Table S17). These results highlight how disease-causing mutations and aging can intersect to fuel proteostatic demise. More generally, our results suggest that the aggregation of increasing numbers of proteins, rather than simply enhanced accrual of specific aggregates, may be an important feature of segmental aging in telomerase deficiencies.

### Age-associated aggregation of proteins linked to disease

These observations led us to ask whether age-dependent protein aggregation involves proteins that are known to be mutated in human diseases. Loss-of-function mutations in many proteins can drive degenerative phenotypes, but we wondered whether aggregation of these proteins in the context of normal aging might underlie slower, progressive tissue defects during normal aging by inactivating them in a non-genetic manner. We focused our analysis on proteins that are genetically linked to diseases as annotated in OMIM (Online Mendelian Inheritance in Man; Table S18). In the heart of old killifish, we observed a substantial increase in the aggregation propensity of A-type nuclear lamins (LMNA) (Figure 7A). Genetic mutations in *LMNA* cause a broad spectrum of diseases, including Hutchinson Gilford Progeria Syndrome (HGPS), muscular dystrophy, and dilated cardiomyopathy (Worman, 2012). Progeria patients experience accelerated aging and early death, often from stroke or coronary artery disease (Eriksson et al., 2003; Merideth et al., 2008).

**Figure 7.**
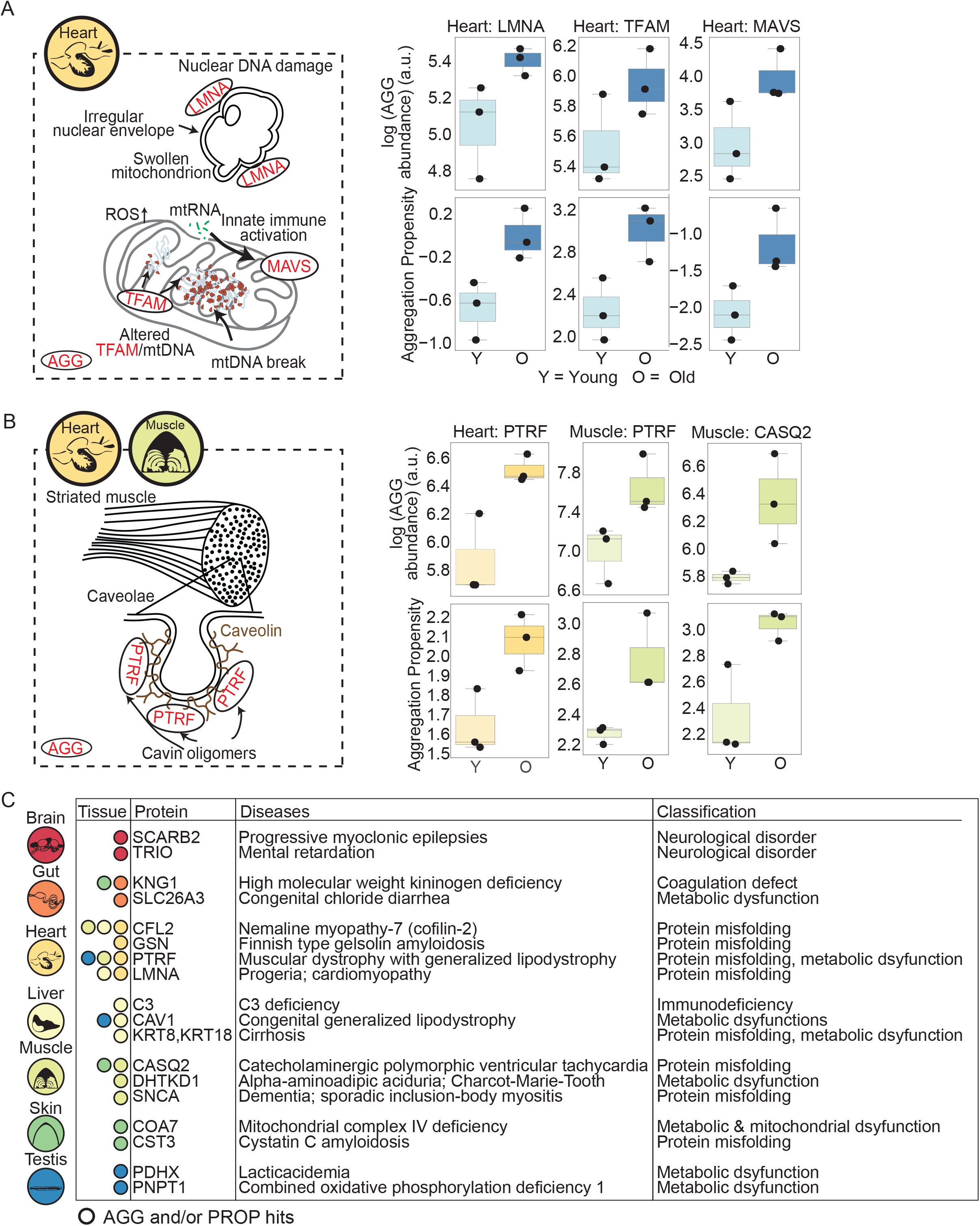
Disease-associated proteins with increased aggregate or aggregation propensity during physiological aging. (A) Cartoon illustration of the functions of proteins with an age-associated increase in aggregate (AGG) and aggregation propensity (PROP) identified in heart and box plot representation of their aggregate (AGG) and aggregation propensity (PROP) levels in young and old animals. Disease-associated proteins and other key proteins involved in mitochondrial unfolded protein response and apoptosis were highlighted here. The box shows the quartiles of the data while the whiskers extend to show the rest of the distribution on protein abundance (y-axis) in old (O) or old *TERT^Δ8/Δ8^* (OT) samples. Each dot represents the AGG abundance or PROP value from an individual fish. (B) Cartoon illustration of disease-associated proteins important for striated muscles function and box plot representation of their aggregate (AGG) and aggregation propensity (PROP) levels in old and young killifish. The selected proteins are known to form oligomeric species and showed an age-associated increase in aggregate and aggregation propensity in two tissues, namely heart and muscle, where striated muscles were isolated in killifish. The box shows the quartiles of the data while the whiskers extend to show the rest of the distribution on protein abundance (y-axis) in old (O) or old *TERT^Δ8/Δ8^* (OT) samples. Each dot represents the AGG abundance or PROP value from an individual fish. (C) Example of proteins with an age-associated increase in aggregate (AGG) and aggregation propensity (PROP) that are also linked to Mendelian diseases. Mutations in these proteins are known to cause or increase the susceptibility to develop the respective diseases. Protein and disease associations are obtained from Online Mendelian Inheritance in Man (OMIM) database. Results are available in Table S17.

Other examples of proteins associated with Mendelian disease that are aggregation-prone include PTRF in the heart, G6PD in the liver, DHTKD1 in muscle (Figures 7B and 7C). Polymerase I and transcript release factor PTRF/Cavin-1 is essential in the biogenesis of caveolae (Hill et al., 2008), and its mutation causes congenital generalized lipodystrophy with myopathy (Ardissone et al., 2013; Dwianingsih et al., 2010; Hayashi et al., 2009; Rajab et al., 2010; Shastry et al., 2010). Mutations in Glucose-6-phosphate dehydrogenase (G6PD), an enzyme that catalyzes the rate-limiting step of the oxidative pentose-phosphate pathway and has a vital role in oxidative stress resistance during aging (Bermudez-Munoz et al., 2020), cause the common genetic enzymopathy worldwide (Cappellini and Fiorelli, 2008). DHTKD1 is part of the mitochondrial 2-oxoglutarate dehydrogenase complex involved in the degradation of several amino acids (Bunik and Degtyarev, 2008). Mutations in this protein lead to the human disease called Charcot-Marie-Tooth Type 2Q, which is associated with progressive atrophy of neurons and muscles. These data raise the possibility that age-associated protein aggregation of wild-type proteins might drive a greater number of age-associated degenerative diseases than is currently appreciated.

## DISCUSSION

We report the first quantitative profiling of protein aggregation during aging across diverse tissues. To do so, we leveraged a new vertebrate model of aging – the African killifish – and innovative genomic tools that have been developed for this organism, enabling comprehensive profiling of protein expression and aggregation across lifespan. The ability to genetically modify the killifish and rapidly conduct aging experiments also allowed us to examine how genetic disease risk and age interact to destabilize protein quality control. Our work significantly expands the growing repertoire of aggregation-prone proteins, beyond past efforts (Becher et al., 2018; David et al., 2010; Kelmer Sacramento et al., 2020; Walther et al., 2015). It not only uncovers previously unknown protein aggregates but also reveals a surprisingly strong tissue-specificity in protein aggregation during normal physiological aging.

This specificity is likely derived from the interaction between the biophysical features intrinsic to the aggregating protein and the specific defects in the protein quality control network that arise in each tissue. We performed unbiased Monte-Carlo simulation and machine learning analysis (support vector machine classifier) to identify defining features of age-dependent protein aggregates. Some features were shared among protein aggregates from multiple tissues. For example, many proteins that aggregated in old animals shared a charge-patterning profile, with more evenly interspersed regions of short charged and hydrophobic sequences, pointing to the importance of local electrostatic interactions in controlling protein aggregation. However, many enriched biophysical features that emerged from our analyses were most strongly evident in specific tissues. For example, age-dependent aggregates in the brain were enriched in glutamine and asparagine residues, which are characteristic of prion-like domains, a concept we explore in the adjoining paper.

Strikingly, these enrichments could often be explained by specific protein quality control factors that appeared in the aggregate fraction of aged animal tissues. For example, DNAJB6, an Hsp40 that chaperones glutamine- and asparagine-rich prion proteins (Thiruvalluvan et al., 2020), aggregated in the aging brain. Likewise, aggregation of TRiC/CCT was evident in the aging liver, where its clients actin and tubulin also showed enhanced age-dependent aggregation with age. In other systems, anecdotal evidence for co-aggregation of specific chaperone and client proteins has been observed (Dickey et al., 2007; Petrucelli et al., 2004). In addition, ordinarily soluble proteins, including chaperones, become SDS-insoluble when mice are genetically engineered to carry a high load of proteotoxic amyloid (Xu et al., 2013). Our data suggest that this effect may be widespread and promote the tissue-selective breakdown of protein homeostasis during natural aging. Indeed, components of the protein life cycle from birth to death: ribosome, chaperones, and proteasome all showed evidence of tissue-specific aggregation. Future work will be required to determine the functional importance of this behavior in aging, and whether such aggregation of proteostasis factors could also be involved in a ‘snowball’ effect, leading to the formation of additional aggregates throughout lifespan.

Exactly how tightly protein aggregation is coupled with aging has long been debated. To investigate this question, we leveraged the power of the killifish to investigate the interaction between a genetic model of telomerase deficiency and age. In telomerase deficiency syndromes, proliferative tissues age more rapidly than usual, but non-proliferative tissues are largely unaffected. Our proteomic maps established that tissues subject to more ‘rapid’ aging in telomerase mutants experienced more protein misfolding events leading to increased aggregate burden. In dividing stem cells, asymmetric inheritance of damaged proteins is tightly linked to longevity and cell fate; long-lived neuroblasts and germline stem cells are rejuvenated, whereas short-lived intestinal stem cells inherit damage (Bufalino et al., 2013). In addition, thermal proteome profiling revealed stabilization of disordered proteins during mitosis (Becher et al., 2018). During aging, protein stabilization afforded by protein quality control can be lost, leading to the accumulation of misfolded proteins in each cell division. This increase in protein damage may manifest earlier in short-lived proliferative tissues simply because they undergo the most divisions. In cancer, a strong positive correlation has been reported between the number of stem cell divisions in the lifetime of a given tissue and its overall cancer risk (Tomasetti and Vogelstein, 2015). It is tempting to draw parallels between the accumulation of damaged proteins and the increase in mutation burden during the lifetime of individual tissues. Together, these observations suggest that aggregation is a core aspect of aging and that the intersection of genetic disease risk with age can drive significant changes in the proteome.

Aging is one of the most significant risk factors for many diseases across all tissues. Our dataset revealed that protein aggregation might also underlie many of these relationships. We identified multiple disease-associated proteins that had increased aggregate load and aggregation propensity during normal aging in all tissues. Most of these proteins had not previously been known to aggregate with age. In fact, some are linked to diseases that have never been attributed to protein misfolding (Boersema et al., 2018). Thus, aggregation of disease-associated proteins themselves during aging likely extends beyond neurodegeneration and could contribute to progressive deterioration of tissues and increased propensity toward diverse diseases during the course of aging. We also reveal an interesting intersection between protein aggregation and diseases with accelerated aging phenotypes: aggregation of nuclear lamina protein LMNA—known genetic factor for progeria—in aging heart and increased aggregation in a telomerase-deficient progeria model. Together, these observations suggest that protein aggregation may contribute to tissue damage and disease in many tissues in addition to the brain, acting as a ‘trigger’ of dysfunction, just as a mutation might in earlier life.

To the best of our knowledge, our study is the first organism-wide quantitative profiling of total protein and aggregation during natural vertebrate aging. We observed aging signatures that are conserved with worms, fruit flies, and mice while also obtaining unique insight into the tissue-specific nature of age-associated aggregation at the protein identity, function, and feature levels. We paired individual total protein and aggregate samples in our analysis which gave us the unique power to identify protein autonomous and tissue intrinsic contributors of protein aggregation in aging. Future work is needed to isolate different cell types from these tissues and understand their differences in protein aggregation profile. Cell-autonomous regulation of proteostasis in aging has previously been investigated in the context of neurodegenerative disease models and aging in worms (Morimoto, 2020; Prahlad and Morimoto, 2011; Taylor and Dillin, 2013). Global proteome remodeling during aging was reported in worms (Walther et al., 2015), but studies in higher eukaryotes revealed more subtle changes (Ori et al., 2015; Walther and Mann, 2011; Yu et al., 2020). Therefore, it is critical to develop tools and genetic models that allow the investigation of tissue crosstalk. Key questions include how damaged proteins are formed, detected, propagated, and degraded. Our observation of Mendelian disease proteins aggregating during aging without underlying mutation begs the question of the origin of misfolding events. One possible explanation lies in protein mistranslation (Gonskikh and Polacek, 2017).

It has been proposed that avoiding protein aggregation has been a driving force in the evolution of protein sequences (Wright et al., 2005). Models invoking this cost have been used to explain the strong anticorrelation between gene expression levels and evolutionary rate; the cost of mutation-induced aggregation is greater for abundant proteins than for rare ones (Drummond and Wilke, 2008). We also observed this anticorrelation relationship between non-synonymous mutation rate (Ka) and total protein and aggregate levels, where it was consistently stronger, across young killifish tissues (Figure S7C). However, in keeping with theory suggesting limited evolutionary selection on aging, the relationship decayed in all tissues from aged animals with one exception: the brain. That is, in contrast to other tissues, age-dependent aggregates in the brain are likely to arise from proteins that have experienced higher rates of diversification (Figure S7D). Of course, this relationship can arise from either drift or selection, and it is impossible to extrapolate with confidence from a single organism. Nonetheless, these observations raise the possibility that innovation of the vertebrate nervous system may be inextricably linked to a key aspect of vertebrate aging.

Age-related diseases, ranging from neurodegeneration to cancer and even COVID, are among the greatest threats to public health in modern society. If effective therapies are not found, they also have the potential to bankrupt the developed and developing world. For many of these conditions, protein aggregation and proteostasis dysfunction are among the major underlying pathologies. Yet efforts distinguishing cause from effect and developing means for intervention are often sparse, and models that faithfully recapitulate vertebrate aging are urgently needed. This study provides a robust platform to begin investigating these questions mechanistically, across tissues, and at a systems level.

## AUTHOR CONTRIBUTIONS

Y.R.C., I.H., A.B., and D.F.J. designed this study with initial help from I.Z. I.H. isolated killifish organs, and Y.R.C. and I.Z. isolated samples for mass spectrometry analysis with the help of I.H. and did silver stain analysis. Y.R.C designed and implemented mass spectrometry data analysis with input from I.H., P.P.S., A.B. and D.F.J. Y.R.C. constructed yeast strains expressing killifish proteins with help from I.H. and B.E.M. and quantified yeast microscopy images. Y.R.C. performed protein feature analysis and built machine learning models. P.P.S. performed GSEA analysis and calculated the evolutionary rate of killifish proteins. P.P.S. and Y.R.C. conducted independent code check on the analysis. E.M. and U.G. performed the FUCCI FACS analysis under the guidance of I.H. Y.R.C., I.H., A.B., and D.F.J. wrote the manuscript, and all the authors commented on the manuscript.

## DECLARATION OF INTERESTS

The authors declare no competing interests.

## ACKNOWLEDGMENTS

We thank Alex Holehouse, Serena Sanulli, and Mardi Bijleveld for critical reading and feedback on the manuscript. We thank the Brunet, Jarosz, and Harel labs, and in particular Olivia Zhou, for stimulating discussion and feedback on the manuscript. We thank Katja Hebestreit and Christine Yeh for their help with formulation of the initial mass spectrometry analyses. We thank Stanford’s mass spectrometry facility, in particular Christopher M. Adams and Ryan Leib, for assistance processing the samples. We thank Parag Mallick and Josh Elias for feedback on mass spectrometry analyses. We thank Kiran Chandrasekher and Ray Futia for independent and blinded quantification of protein aggregates from yeast microscopy images. We thank Sifei Yin for her help with sequence verification of killifish constructs. We thank Susan Murphy (Stanford) and Ashayma Abu-tair (HUJI) for help with killifish maintenance. This work was supported by NIH RF1AG057334 (A.B., D.F.J), R01AG063418 (A.B., D.F.J.), the Stanford Alzheimer’s Disease Research Center and the Zaffaroni Alzheimer’s Disease Translational Program (A.B., D.F.J.), the Stanford Brain Rejuvenation Project (A.B.), the Glenn Foundation for Medical Research (A.B.), the Stanford Systems Biology Seed Grant (I.H., Y.R.C., and I.Z.), Abisch-Frenkel Foundation 19/HU04 (I.H.), Zuckerman Program (I.H.), NIH 1R21AG063739 (I.H.), ISF 2178/19 (I.H.), Israel Ministry of Science 3-17631 and 3-16872 (I.H.), Moore Foundation GBMF9341 (I.H.), BSF-NSF 2020611 (I.H.), and Israel Ministry of Agriculture 12-16-0010 (I.H). I.H. was supported by a Damon Runyon, Rothschild, and Human Frontiers long-term post-doctoral fellowships. Y.R.C. was supported by a Stanford Graduate Fellowship.

## STAR METHODS

Detailed methods are provided in the online version of this paper and include the following.

### Key Resources Table

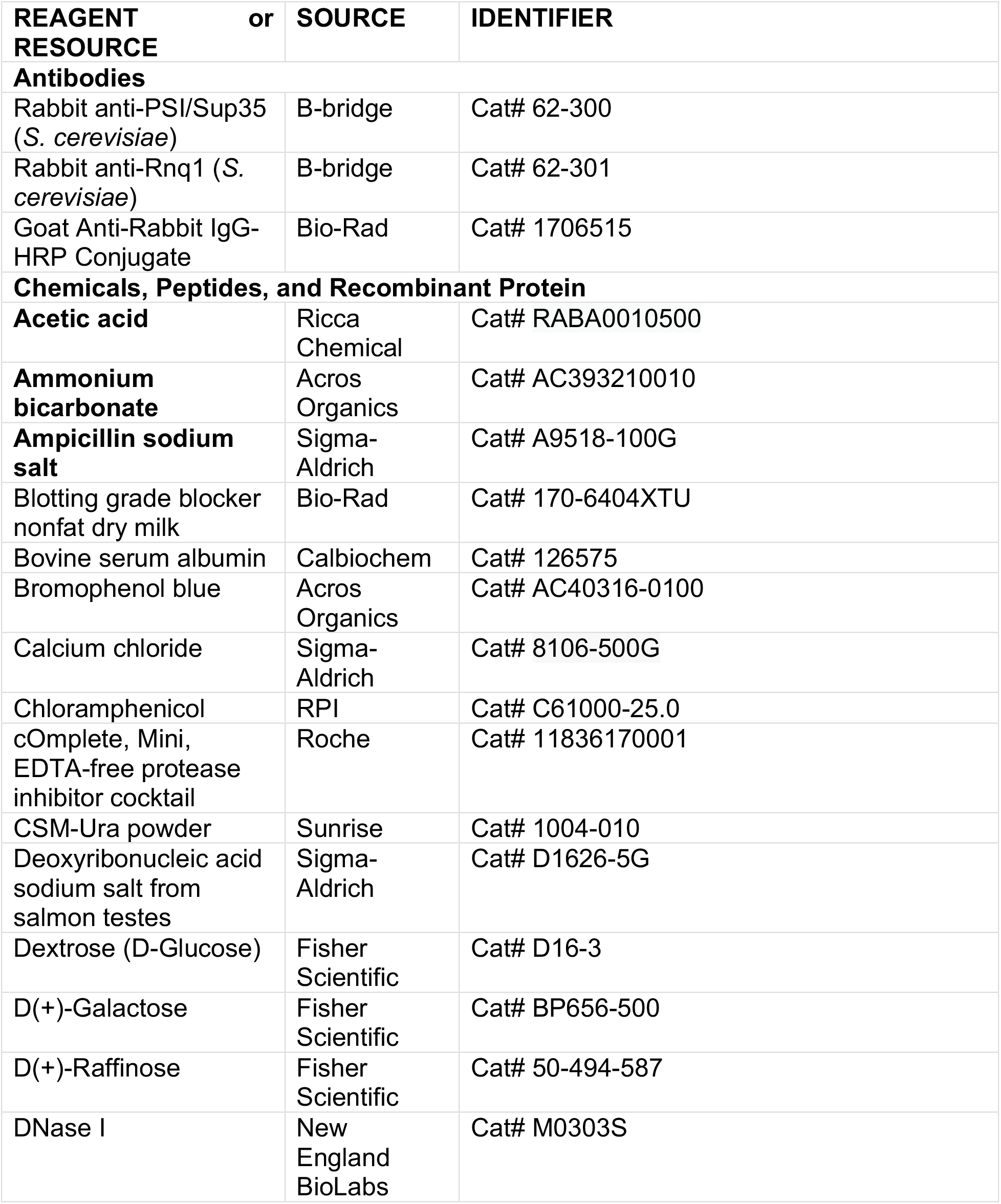
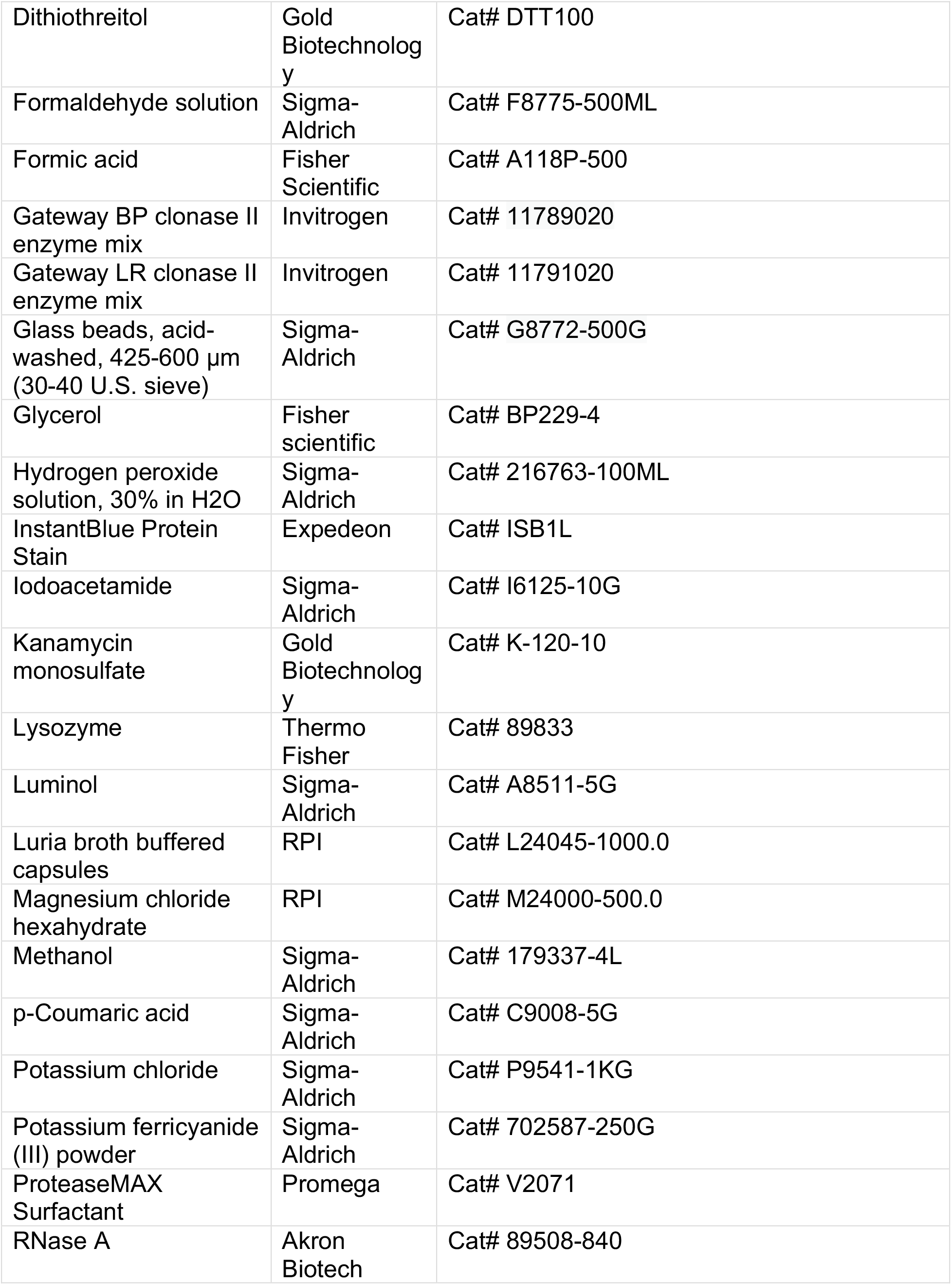
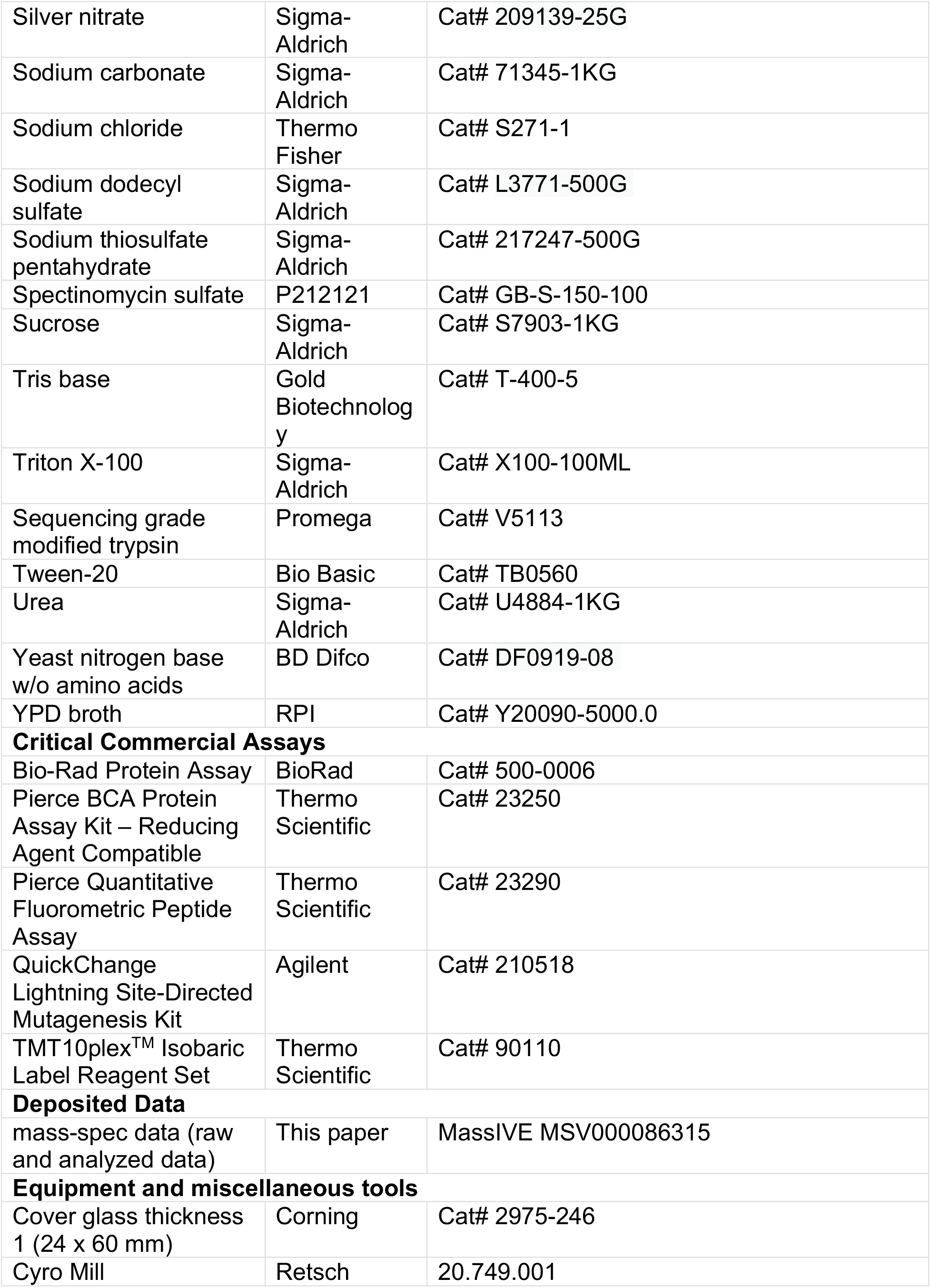
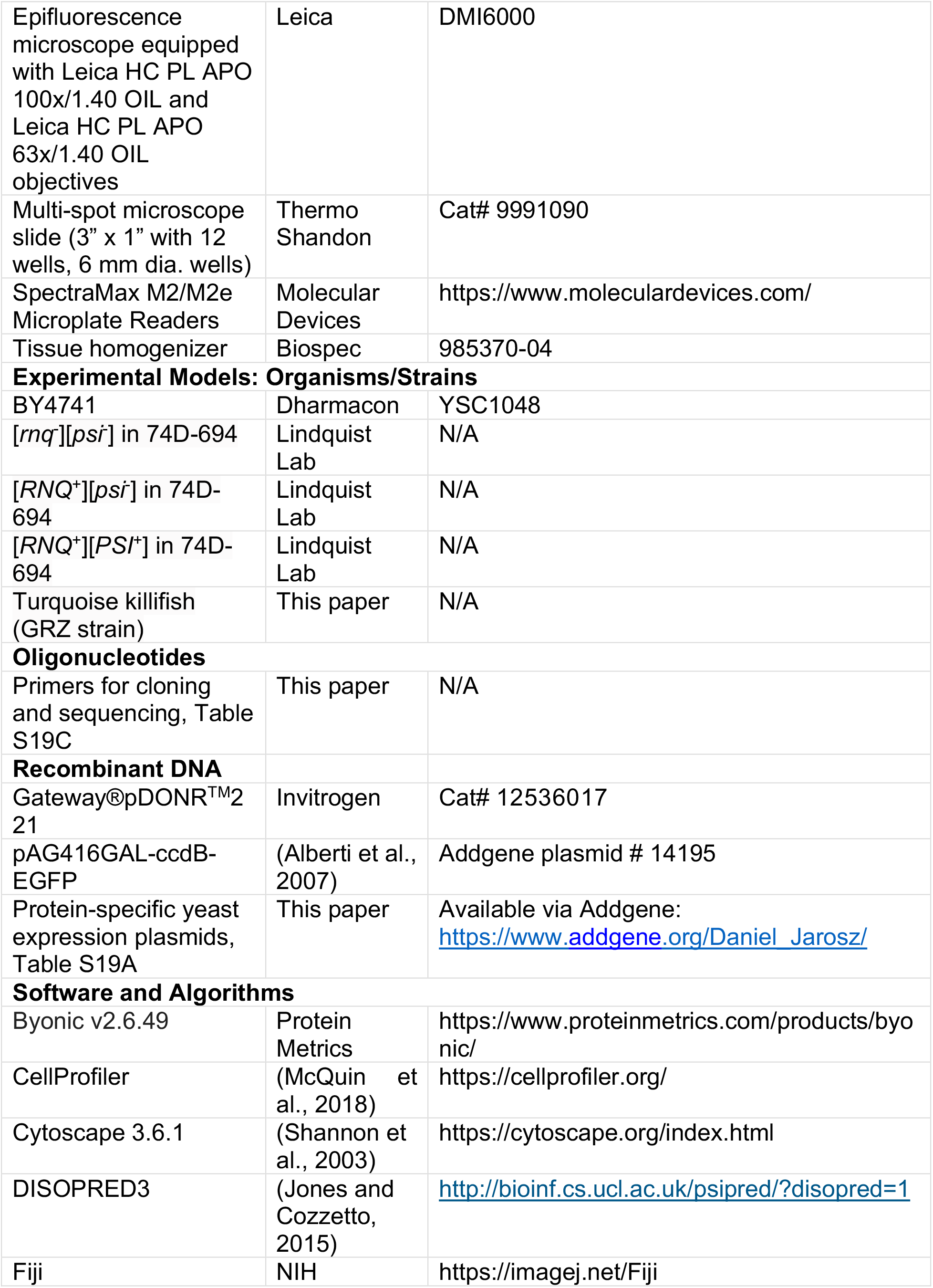
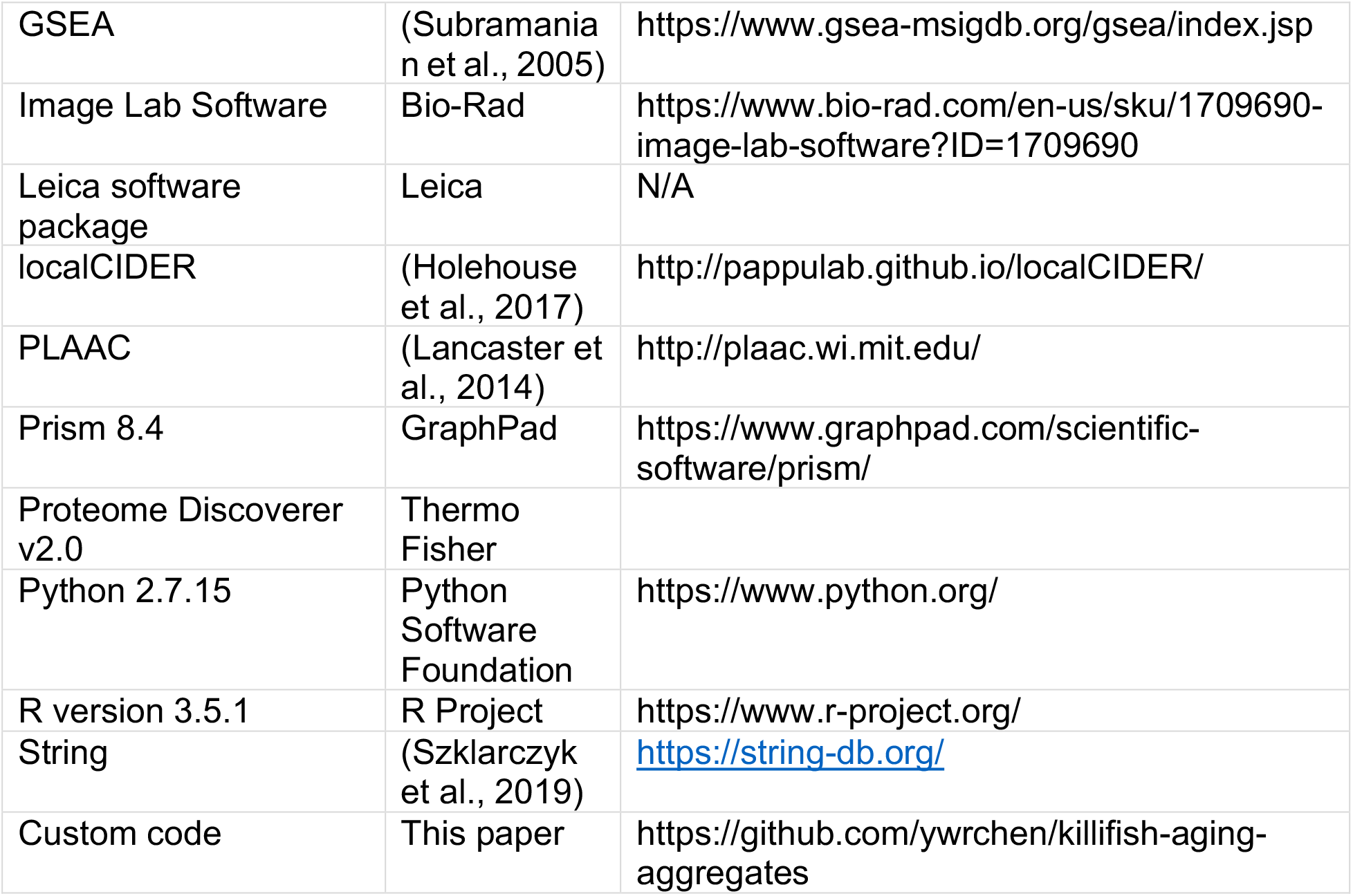

### Resource availability

#### Lead contact and material availability statement

Please contact D.F.J or A.B. for reagents and resources generated in this study.

#### Data and code availability

All raw mass spectrometry reads as well as processed datasets can be found in the MassIVE database (https://massive.ucsd.edu/ProteoSAFe/static/massive.jsp) under ID MSV000086315. The codes and results supporting the current study are available in the Github repository for this paper https://github.com/ywrchen/killifish-aging-aggregates.

### Experimental Model and Subject Details

#### African Turquoise Killifish Strain, Husbandry, and Maintenance

The African turquoise killifish (GRZ strain) were housed as previously described (Harel et al., 2015). Fish were housed at 26°C in a central filtration recirculating system with a 12 hr light/dark cycle (Aquaneering, San Diego) at the Stanford University facility or at the Hebrew University of Jerusalem (Aquazone ltd, Israel). In both facilities, fish were fed twice a day on weekdays and once a day on weekends with Otohime Fish Diet (Reed Mariculture). In these conditions, killifish lifespan was approximately 6-8 months. The *TERT^Δ8/Δ8^* loss-of-function allele (Harel et al., 2015) was maintained as heterozygous (due to fertility issues in homozygous) and propagated by crossing with wild-type fish. All turquoise killifish care and uses were approved by the Subcommittee on Research Animal Care at Stanford University (IACUC protocol #13645) and at the Hebrew University of Jerusalem (IACUC protocol #NS-18-15397-2).

#### Yeast Strain Maintenance

*S. cerevisiae* strains were obtained from the sources indicated (Table S19B). All *S. cerevisiae* strains were stored as glycerol stocks at −80°C. Before use, strains were either revived on YPD (10 g/L yeast extract, 20 g/L dextrose, 20 g/L peptone, sterilized by autoclaving) or on defined medium (2% glucose, 6.7 g/L yeast nitrogen base without amin acids, 20 mg/L histidine, 120 mg/L leucine, 60 mg/L lysine, 20 mg/L arginine, 20 mg/L tryptophan, 20 mg/L tyrosine, 40 mg/L threonine, 20 mg/L methionine, 50 mg/L phenylalanine, 20 mg/L uracil, 20 mg/L adenine, sterilized by autoclaving) as necessary. Antibiotics, or defined drop-out media were used as indicated to maintain plasmid selection. All strains were grown at 30°C unless otherwise indicated.

Yeast strains expressing exogenous killifish proteins were generated by transforming laboratory strain BY4741 (either fresh mid-exponential cells or frozen chemically competent cells) with yeast expression plasmids that encoded proteins of interest. Yeast transformation was carried out using a standard lithium-acetate protocol. First, cells were inoculated into 25 mL of liquid rich medium (YPD) and grown to saturation overnight on a shaker at 200 r.p.m. and 30°C. The cells were then diluted by 25-fold into 500 mL of liquid rich media (YPD) and regrown on a shaker at 200 r.p.m. and 30°C. Once the culture reached mid-exponential phase (OD_600_ ∼0.4 -0.6), the cells were harvested by centrifugation at 2,000 x *g* for 5 min and washed twice in an equal volume of sterile water. The cells were either used directly for yeast transformation or further processed to generate competent cells. To generate chemically competent cells, cell pellets were resuspended in 5 mL of filtered sterile frozen competent cell solution (5% v/v glycerol, 10% v/v DMSO), and 50 µL aliquots were generated in 1.5 mL microcentrifuge tube and stored at −80°C. To ensure good survival rates, aliquots were slowly frozen either using Mr. Frosty freezing container (Thermo Scientific Cat# 5100-0001) or Styrofoam box padded with Styrofoam chips or newspaper (to reduce air space around sample). For yeast transformation, competent cells were thawed in 37°C water bath for 15-30s then centrifuged at 13,000 x *g* for 2min to remove supernatant and resuspended in a transformation master mix (260 μL PEG 3500 50% (w/v), 36 μL 1 M Lithium acetate, 50 μL denatured salmon sperm carrier DNA (2 mg/mL), 14 μL plasmid DNA (0.1-1 μg total plasmid), and sterile water to a final volume of 360 μL). Cells were incubated in the transformation master mix at 42°C for 45 min. Following incubation, cells were harvested, resuspended in 1 mL sterile water, and ∼100 μL was plated on selective medium and incubated at 30°C. Successful transformants typically appeared in 2-3 days and were further propagated in defined liquid drop-out medium (omitted nutrient depending on the plasmid being selected) and stored as glycerol stocks in -80°C.

### Method Details

#### Aggregate Isolation Protocol and Validation with *S. cerevisiae* Strains

We have adapted a standard aggregate isolation protocol (Kryndushkin et al., 2013; Kryndushkin et al., 2017) to separate relatively small oligomeric protein aggregates (Figure S1A) from other membraneless organelles (e.g. stress granules) (Chen et al., 2021). Large membraneless organelles (Jain et al., 2016; Mitchell et al., 2013; Sheth and Parker, 2003) can co-pellet with aggregates if ultracentrifugation steps are performed directly after lysate clarification. Our protocol thus introduces multiple differential centrifugation steps to exclude both large membraneless organelles (e.g. stress granules) (Wheeler et al., 2017) large defined macromolecular complexes (e.g. ribosomes(Ingolia et al., 2012). A detailed comparison of various aggregate isolation protocols follows in the next section on aggregate isolation in killifish.

We first validated this protocol in the widely used yeast laboratory strain BY4741. One liter of BY4741 was grown in rich medium (YPD) until it reached late-exponential phase (OD_600_ ∼1.0) with shaking (250 r.p.m.) at 30°C. The culture was harvested and pelleted at 3000 x *g* for 10 min. The yeast pellet was washed with deionized water twice before proceeding to the aggregate isolation step. We lysed the washed yeast cell pellet in a cryomill (Retsch, for larger cultures over 500 mL) or using acid-washed glass beads (425-600 μm in bead diameter, Sigma-Aldrich, G8772-500G, for smaller culture volumes) in lysis buffer (30 mM Tris-HCl pH 7.5, 40 mM NaCl, 1 mM DTT, 3 mM CaCl_2_, 3 mM MgCl_2_, 5 % glycerol, 1 % triton X-100, EDTA-free protease inhibitor tablets used at the manufacturer’s recommended concentration (Roche cOmplete^TM^ Protease Inhibitor Cocktail, 11836170001)) with 1 mL of lysis buffer per gram of wet cell paste. We then spun the lysate at 800 x *g* for 10 min (spin 1, Eppendorf Centrifuge 5430 with Eppendorf FA-45-30-11 30-spot 45-degree fixed angel rotor) at 4°C to remove cell debris before transferring the supernatants to a new tube and treating with 1 µg/mL RNase A (Akron Biotech, 89508-840), and 20 units/mL DNase (TURBO Dnase, Invitrogen, AM2238) for 30 min on ice. RNase A and DNase treatments were performed to exclude proteins that aggregate exclusively as a result of binding to nucleic acids. Following this incubation, we centrifuged the samples at 10,000 x *g* for 15 min at 4°C (spin 2, Eppendorf Centrifuge 5430 with Eppendorf FA-45-30-11 30-spot 45-degree fixed angel rotor) and kept the supernatant as the whole cell lysate (WCL) fraction (equivalent of tissue lysate (TL) fraction for killifish tissues, see below). A fraction of this WCL was set aside for later analyses. The remainder was loaded on top of a 1 mL 40 % sucrose (in lysis buffer) cushion in an ultracentrifuge tube (Beckman Coulter Ultra-Clear Thinwall Tube, 344057). Higher molecular weight aggregates were then pelleted by ultracentrifugation at 200,000 x *g* for 1 h using Beckman TLS55 swing-bucket rotors (spin 3, 49,000 r.p.m.). We removed the top layers of supernatant carefully and then resuspended the pellet in ∼50 µL of lysis buffer. The resulting aggregate ‘AGG’ fraction was analyzed directly by immunoblot or re-solubilized in 8M urea for further analysis by mass spectrometry.

For the mass spectrometric analysis, 12 µg aggregate (determined by BCA kit, Thermo Scientific Cat# 23250) was re-suspended in SDS-sample buffer and run through approximately 2.5 cm of a 4-15% SDS-PAGE (Mini-PROTEAN TGX Precast Protein Gel, Bio-Rad. Cat #4561086). The top 0.5 cm gel piece near the well (containing the SDS-resistant protein species) as well as the next 2 cm gel piece (containing the SDS-soluble protein species) were excised. Each gel band was reduced with 25 mM DTT, alkylated with 10 mM iodoacetamide, and then digested with trypsin (5 ng/µL) overnight in the presence of 50 mM ammonium bicarbonate. The overnight trypsin digestion was quenched with 5% formic acid in 50% acetonitrile. The digested peptides were recovered from the supernatant and concentrated by speed-vac to remove the acetonitrile solvent, followed by cleaning on a C18 column. The SDS-resistant and SDS-sensitive fractions for each sample were re-constituted in 8 µL and 24 µL of 0.1% formic acid, respectively, and 2 µL of each reconstituted sample was injected onto Orbitrap Fusion Tribrid Mass Spectrometer (Thermo Fisher) for label-free quantification. Mass spectra were analyzed using Proteome Discoverer v2.0 (Thermo Scientific) and the Byonic v2.6.49 search algorithm node for peptide identification and protein inference.

Our data provided high coverage (average and median observed peptide count for a protein was 32 and 6, respectively, and a total of 2600 proteins were identified). Even with this high coverage, abundant ribosomal proteins were depleted in our aggregate fractions relative to expectations from whole cell lysate abundance (Figure S1B), establishing that the aggregate proteome was not simply a sampling of the total cellular proteome (Table S1).

We standardized the protein abundance by calculating the z-score of the total number of spectra for each protein in our yeast aggregate mass spectrometry study and the z-score on protein abundance expressed as mean molecules per cell in the unified *S. cerevisiae* proteome quantification database (Ho et al., 2018) to allow meaningful interpretation of enrichment of specific protein constituents (i.e. processing bodies, stress granules, and ribosomes, Figure S1B). Furthermore, widely used markers of P-bodies (e.g. Edc3 ranked 1933 out of 2600) and stress granules (e.g. Pab1 ranked 119, Pub1 ranked 555, Ded1 ranked 82 out of 2600) were not among the most abundant proteins in the aggregate fractions (Table S2).

To further validate this aggregate isolation protocol (Figure S1A) we used it to examine *S. cerevisiae* strains that harbor specific amyloid aggregates (Rnq1 and Sup35 in [*RNQ*^+^] [*PSI*^+^] strains or Rnq1 in [*RNQ*^+^][*psi^-^*] strains) (Figure S1C). We used the same protocol as described and performed western-blot on the isolated aggregates (pellet in the last ultracentrifugation spin). Specifically, 50 µg aggregate (determined by BCA kit, Thermo Scientific Cat# 23250) from each sample was resuspended in SDS-sample buffer (5X SDS sample buffer: 10% SDS, 50% glycerol, 250 mM Tris-HCl pH 6.8, 10 mM DTT, 0.05% Bromo Phenol Blue) and then split in equal volumes where one sample was boiled (5 min at 95°C) while the other was left unboiled. Both the boiled and unboiled samples were resolved on 12% SDS-PAGE gels (Mini-PROTEAN TGX Precast Protein Gel, Bio-Rad. Cat #4561045). The gel was then transferred onto a 0.2 µm PVDF membrane using the pre-programmed high MW transfer protocol (constant 2.5 A for 10 min) in a Bio-Rad Trans-blot Turbo Transfer System (Cat# 1704150). After transfer, the membrane was submerged in 20 mL of blocking buffer (TBS^T^ + 5% dry milk) for 1 h on a rocker at room temperature. After blocking, the membrane was washed briefly in TBS^T^ twice and incubated with 10 mL of the primary antibody diluted in TBS^T^ (for Rnq1, we used B-bridge rabbit anti-Rnq1 antibody, Cat# 62-301 at a 1:3,000 dilution; for Sup35, we used B-bridge rabbit anti-PSI/Sup35 antibody Cat# 62-300 at a 1:1,000 dilution) on a rocker overnight at 4°C. After the primary antibody incubation, the membrane was washed with TBS^T^ 3 times for 7 min each before incubating with the secondary antibody (goat anti-rabbit IgG-HRP conjugate Cat# 1706515 at a 1:5,000 dilution) for 1hr at room temperature. The membrane was then washed 3 times with TBS^T^ for 5 min each, then 3 times with 0.1 M Tris-HCl pH 8.5 for 5 min each, followed by chemiluminescent detection. The chemo-luminescent reaction was initiated immediately before detection at room temperature by incubating the membrane with 10 mL of solution A (10 mL 0.1 M Tris-HCl, pH 8.5, 100 µL of 44 mg/mL Luminol in DMSO and 42 µL of 14.7 mg/mL p-coumaric acid in DMSO) and 10 mL of solution B (10 mL 0.1 M Tris-HCl, pH 8.5 and 5.5 µL of 30% w/w hydrogen peroxide solution) for 1 min at room temperature.

The amyloid (SDS-resistant) form of Rnq1 was clearly detected in the [*RNQ*^+^][*psi^-^*] strain using this protocol. In the unboiled sample, the Rnq1 antibody reacted with proteins stuck near the top of the well, where SDS-resistant amyloids accumulate; little signal was present at 43 kDa (the molecular weight of soluble Rnq1). After boiling, which re-solubilizes Rnq1 amyloids, we observed strong signal at 43 kDa and depletion of signal in the well.

Similarly, in [*RNQ*^+^][*PSI^+^*] strains we detected the amyloid forms of both Rnq1 and Sup35 in the SDS-resistant fraction (near the top of the well). After boiling, we observed strong signal enhancement for both proteins at their respective molecular weights along with concomitant depletion of signal near the well in the boiled sample (Figure S1C). Thus, our protocol can isolate aggregating proteins including amyloid proteins in native condition (Rnq1 in [*RNQ*^+^][*psi^-^*] and Rnq1 and Sup35 in [*RNQ*^+^][*PSI*^+^]).

#### Isolation of Tissue lysate (TL) and Aggregates (AGG) from Killifish Tissues

Brain, gut, heart, liver, muscle, and skin of 3 young (3.5 months), 3 old (7 months), and 3 old *TERT^Δ8/Δ8^* mutant (Harel et al., 2015) (7 months) male fish were collected at the same time and snap frozen in liquid nitrogen. Because *TERT^Δ8/Δ8^* mutant has testis defects (Harel et al., 2015), only testis from 3 young and 3 old male fish were collected and snap frozen in liquid nitrogen. All procedures were conducted at 4°C unless stated otherwise. Each organ was homogenized using a tissue homogenizer in 100 µL of buffer A (30 mM Tris-Cl pH = 7.5, 1 mM DTT, 40 mM NaCl, 3 mM CaCl_2_, 3 mM MgCl_2_, 5% glycerol, 1% triton X-100, protease inhibitor cocktail tablet used at 1x the manufacturer recommended concentration (Roche cOmplete^TM^ EDTA-free Protease Inhibitor Cocktail, Cat# 11697498001). Homogenization was performed in round-bottom tube (2 mL corning cryogenic vials) to ease lysis. The resulting sample was transferred to 1.5 mL Eppendorf tube for the first centrifugation spin. Lysate was spun at 800 x *g* for 10 min (spin 1) to remove cell debris (Eppendorf Centrifuge 5430 with Eppendorf FA-45-30-11 30-spot 45-degree fixed angel rotor). Supernatants were transferred to a new Eppendorf tube and treated with 100 µg/mL RNase A (Akron Biotech, 89508-840), and 100 µg/mL DNase I (New England Bio, Cat# M0303S) for 30 min on ice. Samples were spun at 10,000xg for 15 min (spin 2 in the same Eppendorf FA-45-30-11 rotor) and the resulting supernatant is the tissue lysate (TL) fraction. A 25 µL aliquot of the TL was kept in a separate tube for protein quantification and mass spectrometry analysis. For isolation of the aggregate (AGG) fraction, all the remaining TL was loaded onto the top of a 1 mL 40% sucrose pad and an additional ∼750 µL (adjusted to balance all ultra-centrifuge tubes) of buffer A was layered on the top in ultra-centrifugation tube (Beckman Coulter Ultra-Clear centrifuge tubes, thinwall, 2.2 mL, 11 x 34 mm, Cat# 347356). The samples were separated by ultracentrifugation for 1 h at 200,000 x *g* (spin 3 at 49,000 r,p.m. in Beckman TLS-55 rotor). The top layers of supernatants were removed, leaving 15-20 µL of liquid at the bottom around the pellet. An additional 30 µL of buffer A was added to rigorously re-suspend these pellets. Protein concentration for TL and AGG samples was assessed by BCA assay (Pierce BCA Protein Assay Kit – Reducing Agent Compatible, Cat# 23250).

Our aggregate isolation protocol should theoretically physically separate protein aggregates (size ranging from ∼ 164–8804 Svedberg units, ∼132 – 968 nm in diameter or ∼1.03 x 10^6^ – 4.03 x 10^8^ kDa assuming a spherical shape, see theoretical calculation below) from the soluble proteome, large protein complex (e.g. ribosome in 40-80 Svedberg units, ∼ 20-30 nm in diameter or 4.5 MDa; spliceosome with size ranging from ∼30-100 Svedberg units, ∼30 nm in diameter, or ∼2-20 MDa (Spann et al., 1989; Will and Luhrmann, 2011; Zhang et al., 2017)), subcellular organelles, and large biomolecular condensates (e.g. P-bodies). There are two main differences between our aggregate isolation protocol and those used in some other studies of age-dependent protein aggregation (David et al., 2010; Kelmer Sacramento et al., 2020; Walther et al., 2015). First, our protocol identifies both SDS-soluble and SDS-resistant aggregates in native conditions (the entire isolation is performed at 4 °C and we omitted EDTA to preserve the stability and function of metal-dependent proteins). Second, our protocol enriches oligomeric aggregates that are bigger than large protein complexes but smaller than membraneless organelles (a detailed calculation of aggregate size is described in the theoretical calculation section below).

In the David et al. *C. elegans* study (David et al., 2010; Lechler et al., 2017), aggregates were isolated that remained insoluble in 0.5% SDS (pellet fraction) in RIPA buffer (50 mM Tris pH 8, 150 mM NaCl, 5 mM EDTA, 0.5% SDS, 0.5% SDO, 1% NP-40, 1 mM PMSF, Roche Complete Inhibitors 1×) after 3 rounds of 20,000 x *g* centrifugation for 20 min at 4 °C. Aggregates isolated by this protocol would be expected to be larger than 1650 S (>419 nm in diameter, based on Eppendorf F-45-30-11 rotor with k-factor of 508 at a maximum speed of 20,817 x *g*). Another *C. elegans* study from Walther et al. (Walther et al., 2015) analyzed insoluble proteins after a brief 1 min spin at 1,000 x *g* for lysate clarification (in 50 mM Tris-HCl pH 8.0, 0.5 M NaCl, 4 mM EDTA, 1% (v/v) Igepal CA630, cOmplete^TM^ proteinase inhibitor cocktail) followed by ultracentrifugation at 500,000 *g* x 10 min. All steps were carried out at 4 °C. The aggregate size is expected to be larger than 268 S (>169 nm in diameter, calculated based on using Eppendorf F-45-30-11 for lysate clarification step and Beckman Type 70 Ti fixed angel rotor (k-factor is 44 at a maximum speed 504,000 x *g*) for ultracentrifugation). Thus, the two *C. elegans* studies identified aggregates that are either large (Walther et al., 2,799 proteins larger than 268 S) or both very large and SDS-resistant (David et al., 698 proteins larger than 1650 S). There are also some differences in the thresholds that were applied to identify positive hits between these studies (Figure S1G; Table S3A-B; Table S3E-F).

In the vertebrate (killifish and mice) brain study from Kelmer Sacramento et al. (Kelmer Sacramento et al., 2020), the initial lysis buffer (4% SDS, 100 mM HEPES, pH 8, 1 mM EDTA, 100 mM DTT) contains 4% SDS, which can disrupt the native conformation of oligomeric aggregates species. The aggregates examined in this study were pellets collected after two ultracentrifugation spins at 100,000 x *g* for 30 min at 20 °C where the input was supernatant (brain lysate) from an initial 20,000 x *g* spin for 5 min. The isolated 4% SDS-denatured aggregates are expected to be 670 - 6600 S (268∼837 nm in diameter, calculated based on using Eppendorf F-45-30-11 rotor for the lysate clarification step and Beckman TLS-55 swinging-bucket rotor for the ultracentrifugation (k-factor of 50 at a maximum speed of 259,000 x *g* in aqueous solvent). In this study, 964 and 74 proteins were identified as aggregates in old mice and killifish brains, respectively (young animals were not analyzed; Figure S1G; Table S3D-E).

Finally, in mammalian stress granule purifications (Jain et al., 2016), the cells were first lysed with lysis buffer (50 mM Tris HCl pH 7.4, 100 mM potassium acetate, 2 mM magnesium acetate, 0.5 mM DTT, 50µg/ml heparin, 0.5% NP40, 1:5,000 antifoam emulsion, 1 complete mini EDTA free protease inhibitor tablet/ 50ml of lysis buffer) on ice. Lysates were further treated with an initial centrifugation step at 1,000 x *g* for 5 min, a second centrifugation at 18,000 x *g* for 20 min (input was the supernatant from the first spin), and a final centrifugation step at 850 x *g* for 2 min at 4°C (input was the pellet fraction from the second spin). The stress granule cores were further enriched through affinity purification (immunoprecipitation with specific antibodies to query proteins such as G3BP) of supernatant from the last 850 x *g* spin. The stress granule core is estimated to be larger than 2038 S (> 465 nm in diameter).

Thus, our protocol, which discards pellets in our spin 1 (800 x *g* for 10 min) and spin 2 (10,000 x *g* for 15 min), should deplete for stress granules whereas the protocols used in *C. elegan*s studies (David et al., 2010; Walther et al., 2015) would enrich them. Because the aged killifish and mouse brain experiments were performed at 20°C with 4% SDS in the lysis buffer (Kelmer Sacramento et al., 2020), any SDS resistant particles between 268-837 nm in diameter would also be isolated as aggregates. By contrast, our protocol captures both SDS-soluble and SDS-resistant aggregates while excluding large membraneless organelles and defined macromolecular assemblies (Jain et al., 2016; Mitchell et al., 2013; Sheth and Parker, 2003).

#### Silver Stain on Killifish Samples

Equal amounts (1 µg) of tissue lysate (TL) and aggregate (AGG) fraction from brain and liver were used for silver stain analysis. The samples were resuspended in SDS-sample buffer (5X SDS sample buffer: 10% SDS, 50% glycerol, 250 mM Tris-HCl pH 6.8, 10 mM DTT, 0.05% Bromophenol Blue) without boiling and resolved on 12% SDS-PAGE gels. The gels were then fixed in 40% methanol/10% acetic acid for one hour at room temperature. Next, the gels were incubated in 100 mL of freshly made ‘yellow mix’ (5 g of K_3_Fe(CN)_6_ and 8 g of Na_2_S_2_O_3_·5H_2_O grinded to powder and dissolved in 150 mL water) for 5 min and subsequently rinsed in dH_2_O until they became colorless. The gels were then incubated in 30 mL of 120 mM silver nitrate (4.1g AgNO_3_ in 200 mL of H_2_O, store in dark at 4°C) for 30 min followed by a brief rinse with 2-3X volume of H_2_O and another brief rinse with 2.9% Na_2_CO_3_. Finally, the gels were incubated with 50 mL of freshly made developer (2.9% Na_2_CO_3_ pre-heated to ∼ 60°C and 100 µL of fresh 37% formaldehyde solution Sigma Cat# F8775). Once the desired band intensity was achieved, the reaction was quenched by immediately adding 5% acetic acid to the developer solution and quickly rinsing off all solution with water. The silver-stained gel was imaged on Bio-Rad Gel Doc EZ Gel Documentation System (Cat# 1708270).

#### Theoretical Calculation of the Size Distribution of High Molecular Aggregate

We performed theoretical calculations of the size distribution of high molecular moieties (including aggregates), based on the parameters of our protocol. Because they are based on fundamental physical principles, such calculations should be independent of the sample origin (i.e. yeast, killifish, or other).

Particles with a size larger than 2.06 x 10^6^ S (Svedberg units) should pellet during the 10 min spin at 800 g (spin 1). Particle with a size larger than 8804 S should pellet during the 15 min spin at 10,000 x *g* (spin 2). Particles with a size larger than 164 S should pellet during the 1 h spin at 200,000 x *g* (spin 3). Therefore, the size of the high molecular weight aggregate should range from 164 S to 8804 S. Calculation for each spin is as follows:

Input samples for spin 1 and spin 2 are prepared in 1.5 mL Eppendorf tubes and separated through centrifugation with Eppendorf FA-45-30-11 rotor (max radius 10.1 cm, min radius 8.9 cm, 30-position fixed 45-degree angle rotor that works with Eppendorf Centrifuges 5430R). The FA-45-30-11 fixed angle rotor has a k-factor of 508 (a measure of the rotor’s pelleting efficiency, in S·h units where S is the Svedberg unit) at a maximum speed of 14,000 r.p.m. and a maximum rcf of 20,817 x *g* (details available at http://www.biocenter.hu/pdf/Eppendorf10.pdf).

At 800 x *g*, the adjusted k-factor is:

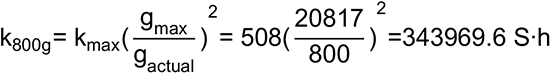

At 10,000 x *g*, the adjusted k-factor is:

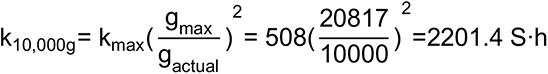

At 20,000 x *g*, the adjusted k-factor is:

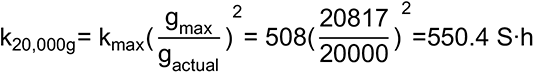

The k-factor for the swinging bucket rotor was calculated as follows based on the maximum and minimum radius of the rotor (r_max_ and r_min_) and the centrifugation speed (in r.p.m.):

Rotor TLS-55 (r_max_ = 76.5 mm, r_min_ = 42.2 mm, k-factor at maximum speed 55,000 r.p.m. is 50 S·h in water at 20°C and 130 S·h in 5-20% sucrose gradient at 5°C, https://btiscience.org/wp-content/uploads/2014/04/TLS55.pdf) that is compatible with Beckman Coulter Optima MAX-TL table-top ultracentrifuge spin at 49,000 r.p.m. (200, 000 x *g*) has a k-factor of:

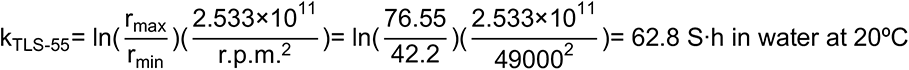

Rotor SW50.1Ti (r_max_ = 107.3 mm, r_min_ = 59.7 mm, k-factor at maximum speed 50,000 r.p.m. is 59) that is compatible with Beckman Coulter Optima MAX-TL table-top ultracentrifuge spin at 40,800 r.p.m. (200,000 x *g*) has a k-factor of:

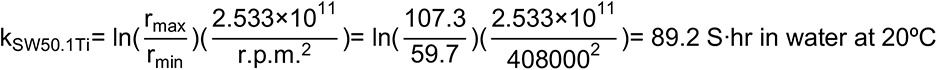

Spin 1: After a 10 min spin at 800 x *g* using Eppendorf centrifuge 5430R, the minimum size of the pellet is

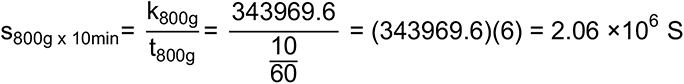

Spin 2: After a 15 min spin at 10,000 x *g* using Eppendorf centrifuge 5430R, the minimum size of the pellet is:

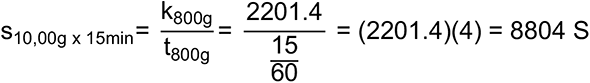

Spin 3: Temperature, density of solution, and speed all affect the pelleting efficiency (or k-factor) so we extrapolated the adjusted k-factor from condition that is closest to ours. Given that the k-factors in established run conditions with 5-20% sucrose at maximum speed at 5 °C is 130 S·h (the particle density is 1.3 g/mL) and 50 S·h in water at maximum speed at 20 °C, we estimated the adjusted k-factor based on the prior condition (our ultracentrifugation is done in 40% sucrose for 1 h at 200,000 x *g* at 4 °C).

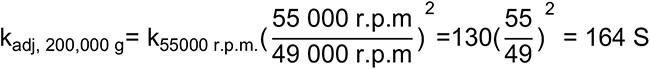

Because 40% sucrose is denser than 5-20% sucrose, it is reasonable to assume that actual pelleting efficiency in our condition is even higher (the same particle will travel more slowly in denser solution) resulting in aggregates with sedimentation coefficient larger than 164 S after 1 hour of ultracentrifugation.

We assumed the aggregate is a perfect sphere and estimated the molecular weight and diameter of the particles based on the following equations.

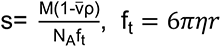

where s is the sedimentation coefficient (1S = 10^-13^ seconds), M is the molecular weight, v̄ is the partial specific volume (for generic protein v̄ is roughly 0.73ml/g), ρ is density of the solution (density of 40% sucrose is 1.176g/mL at 20°C, assuming the density doesn’t change drastically at 4°C), f_t_ is the frictional coefficient, N_A_ is the Avogadro’s number, r is the radius, and *η* is viscosity (viscosity of 40% sucrose is 11.44cP or 11.44 g/(m·s) at 5°C).

The radius of the aggregate can be estimated as

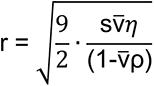

and the molecular weight can be subsequently estimated as

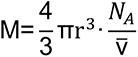

Substituting the particle sedimentation coefficient of 164 S and 8804 S respectively into the equations, we arrived at a size estimate of 66 – 484 nm (radius) and 1.03 x 10^6^ – 4.03 x 10^8^ kDa for the isolated aggregates.

In summary, our aggregate isolation protocol should theoretically physically separate protein aggregates (size ranging from ∼ 164–8804 Svedberg units, ∼132 – 968 nm in diameter or ∼1.03 x 10^6^ – 4.03 x 10^8^ kDa assuming spherical shape) from the soluble proteome, large complex (e.g. ribonucleoprotein particles spliceosome with size ranging from ∼30-100 Svedberg units, ∼30 nm in diameter or ∼2-20 mega-Dalton (Spann et al., 1989; Will and Luhrmann, 2011; Zhang et al., 2017), subcellular organelles, and large biomolecular condensates (e.g. P-bodies) (Figure S1D).

#### Mass Spectrometry Sample Preparation and Analysis

Following tissue lysate (TL) extraction and aggregate (AGG) isolation, samples were re-suspended in 8M urea and ProteaseMAX (Promega) and were subsequently subjected to reduction (with 10 mM DTT for 30 min at 55°C) and alkylation (with 30 mM acrylamide for 30 min at room temperature) followed by trypsin digestion (1:50 concentration ratio of sequencing grade trypsin to total protein overnight at 37°C followed by quenching with 25 µL of 50% formic acid to pH below 3.0). The digested peptides from different samples of the same organ were separately quantified and equal amount of peptide samples were labeled with TMT10plex mass tag before mass spectrometry analysis. In particular, equal amounts of TL or AGG (3-10 µg of peptide depending on the organ/tissue) from each sample was labeled with 9 different TMT-10plex tags accordingly to the manufacturer’s protocols (cat# 90110, Thermo Scientific). The same mass tag and sample assignment was maintained throughout the entire study (old samples were labeled with TMT^10^-126, TMT^10^-127N, and TMT^10^-127C respectively; young samples were labeled with TMT^10^- 128N, TMT^10^-128C, and TMT^10^-129N respectively; and *TERT^Δ8/Δ8^* samples were labeled with TMT^10^-129C, TMT^10^-130N, and TMT^10^-130C respectively). One ninth of each sample (equal amount of peptide across all TL or AGG samples from a tissue/organ) were pooled after trypsin digestion and labeled with the 10^th^ TMT tag to serve as the reference channel for internal normalization. Post-labeling, each set of samples were further cleaned up with C18 peptide desalting columns and went through high pH reverse phase fractionation into 3 (brain, liver, and gut) or 4 (heart, muscle, skin, and testis) fractions and all fractions were run independently on an Orbitrap Fusion (Thermo Scientific) mass spectrometer coupled to an Acquity M-Class nanoLC (Waters Corporation). Data searches were conducted with killifish proteome downloaded from NCBI release 100 (available in the MassIVE dataset and the GitHub repository for this paper). Mass spectra were analyzed using Proteome Discoverer v2.0 (Thermo Scientific) for MS3 quantification of tandem mass tag reporter ions and the Byonic v2.6.49 search algorithm node for peptide identification and protein inference. Briefly, a mass spectrometry analysis allowed for fully tryptic digestion with up to two missed cleavages. A 12 ppm mass accuracy was tolerated for precursor and MS3 HCD fragments, *i.e.*, reporter ions, and 0.3 Da mass accuracy for CID fragmentation at the MS2 level. Static modifications include cysteine carbamidomethylation and TMT labels on peptide N-termini and lysine residues. Oxidation of methionine and deamidation of aspartate and glutamine were considered as dynamic modifications. Peptides and proteins were cut at the 1% FDR level using the Byonic node. Reporter ion intensities were normalized against a pooled sample containing each of the other samples in a given sample run and reported relative to these pooled samples. These ratios were exported for analysis at both the protein and PSM (Peptide Spectrum Match) level. The mass spectrometry raw data, summary table of the reporter ion ratios at protein and PSM levels, as well as protein sequence FASTA files use for the search have been deposited to MassIVE with a dataset id as MSV000086315.

#### Annotation of the NCBI Genes Models of Turquoise Killifish

We used the African turquoise killifish (*N. furzeri*) NCBI annotation release 100 for our analysis. The majority of the gene models in this annotation only have a locus number. Therefore, we re-annotated all the killifish gene model names based on orthology analyses with 40 species including mammals, fish and invertebrates. We selected the consensus symbols for the locus assigned by NCBI as our final symbols and re-annotated the genes using a naming scheme Gene_Name(n of m) if there were duplicates in the killifish genome. For most of the analyses, human orthologs were reported or used in database search, unless otherwise noted. The annotation and human orthologs used in this killifish study can be found in Table S2C.

#### Mass Spectrometry Data Normalization and Analysis of Age-associated Changes

The target protein results including the reporter ion ratios and total number of spectra assigned to peptides from this protein (# PSMs) were further processed to infer the abundance of each protein in each sample. First, the human contaminants were removed. Next, the protein abundance of each sample was inferred from its PSM contribution (or equivalently each TMT10plex tag), calculated by multiplying the total number of PSMs for a protein by the fraction of reporter ion signal that came from this channel (ratio of query channel divided by sum of the ratios across all channels, note that the TMT-131 was the normalization channel and contributed as 1 to the overall signal). Because equal amounts (by mass) of peptides were loaded in each channel, we normalized the sum of PSMs for all proteins in a channel to a constant of 100,000. The resulting normalized counts of PSMs for a protein in a sample represent the final reported protein abundance (PSMsNorm). We log2-transformed the protein abundance (log2_PSMsNorm) and the resulting protein abundance for each tissue effectively followed a normal distribution (Figure S3A). We then used the normally distributed log2-transformed protein abundance to perform parametric statistical tests (i.e. Student’s t-test), as is often done for proteomics datasets (Klann et al., 2020; Li et al., 2019; Mirzaei et al., 2017; Navarrete-Perea et al., 2018; Nusinow et al., 2020; Zhang and Elias, 2017).

The age-associated changes in a protein in either tissue lysate or aggregate fraction were calculated as the fold change in the average abundance of a protein between the two age/disease groups. Both fold change (i.e. OvY_FC for the fold change of old divided by young) and log2-transformed fold change (i.e. OvY_logFC was the log2-transformed fold change of old divided by young) are reported in Table S2. The p-values were assessed using a Student’s t-test with log2-transformed protein abundance of the two age/disease group (i.e. OvY_pval).

We defined the term ‘aggregation propensity’ (PROP) to infer the intrinsic likelihood of a protein to aggregate, scored by dividing the abundance of a protein in the aggregate fraction (AGG) by its tissue lysate (TL) abundance. Note that this metric can only be reported when a protein is identified in both the TL and AGG fractions. Proteins that were only detected in AGG but not in TL make up about 0.8-1.7% (median 1.3%, average 1.2%) of total AGG signal and their abundance changes were analyzed at the AGG level only. Because each channel represents the tissue sample from an individual fish, we reported the aggregation propensity of a protein for each sample. The age-associated changes in aggregation propensity (i.e. OvY_prop_FC) were calculated as the fold change in the average aggregation propensity of a protein between the two age/disease groups. The log2-transformed fold change (i.e. OvY_prop_logFC) was reported as well. Student’s t-tests were performed on the log2-transformed aggregation propensity of the different conditions (young, old, old *TERT^Δ8/Δ8^* mutants). The resulting p-values were reported (OvY_prop_pval) to assess whether the changes between conditions were statistically significant.

#### Reproducibility Between Proteomic Samples

The reproducibility of the proteomic datasets was assessed by comparing the measured protein abundance between one biological replicate and another (log2-transformed normalized PSMs, log2_PSMsNorm). There are 3 biological replicates for each condition (young, old, old *TERT^Δ8/Δ8^* mutants), and they were all compared to one another both for tissue lysate (TL) and aggregates (AGG). The resulting Pearson’s correlation coefficient was reported (Figure 1D and Figure S1F).

#### Biophysical and Sequences Features of Aggregation-Prone Proteins

The complete list of properties analyzed include:

- “AA Length” – the protein amino acid length;
- “count Neg” –the number of negatively charged residues (D, E);
- “count Neut” – the number of polar neutral residues (S, T, N, Q);
- “count Pos” – the number of positively charged residues (H, K, R);
- “delta” – protein charge patterning parameter obtained when calculating kappa using the localCIDER developed by the Pappu lab (Holehouse et al., 2017)
- “deltaMax” – the maximum possible delta value for a sequence of this composition when calculating kappa (Das and Pappu, 2013);
- “kappa” – protein charge patterning parameter defined as a ratio of the sequence’s delta over the maximum possible value for a sequence of that composition (Das and Pappu, 2013);
- “kappa_afterPhos” – the protein charge patterning parameter kappa assuming full phosphorylation (Das and Pappu, 2013);
- “Omega” –Omega defines the patterning between charged/proline residues and all other residues (Martin et al., 2016);
- “Disordered Fraction” – the fraction of total residues with a DISOPRED3 score above 0.5 (Jones and Cozzetto, 2015);
- “DISOPRED max disorder” – the maximum stretch of disorder predicted by DISOPRED3 (Jones and Cozzetto, 2015);
- “FImaxrun” or “FoldIndex max disorder” – the maximum stretch of disorder predicted by FoldIndex (Prilusky et al., 2005);
- “FImeancombo” – the disorder score for (disorder score for entire protein defined as 2.785<H> - |<R>| - 1.151, where <H> is the Uversky hydropathy score and <R> is mean charge;
- “FFInumaa” – the number of amino acids predicted to be disordered by FoldIndex (Prilusky et al., 2005);
- “FoldIndex disorder fraction” – the fraction of total disorder residues predicted by FoldIndex;
- “Frac_aliphatic” – the fraction of non-polar aliphatic residues (A, V, L, I, and M);
- “Frac_aromatic” – the fraction of non-polar aromatic residues (F, Y, and W); “Frac_Neu” – the fraction of polar neutral residues (S, T, N and Q);
- “Frac_Neg” – the fraction of negatively charged residues (D and E);
- “Frac_pos” – the fraction of positively charged residues (H, K, and R);
- “FracCharged” – fraction of charged residues (H, K, R, D, and E);
- “fraction of chain expansion” – the fraction of residues that contribute to chain expansion (E/D/R/K/P) (Holehouse et al., 2017);
- “fraction of disorder promoting” – fraction of residues that is predicted to be ‘disorder promoting’ in TOP-IDP-scale (Campen et al., 2008);
- “Frac_QN” – the fraction of Q and N residues;
- “MeanNetCharge” – absolute mean net charge of a particular protein sequence;
- “MW” – the protein molecular weight;
- “Michelitsch-Weissman score” – prion score predicted by the method developed by the Weissman lab, equivalent to maximum number of Qs and Ns in a window of at most 80 amino acids (Michelitsch and Weissman, 2000);
- “MWlen” – the length of prion-like region predicted from the method developed by the Weissman lab (Michelitsch and Weissman, 2000);
- “PAPAprop” or “PAPA prion propensity” – the predicted prion propensity by PAPA (Toombs et al., 2012);
- “NCPR” – the net charge per residue of a protein;
- “NLLR” or “normalized PLAAC score” –the normalized prion score NLLR predicted by PLAAC (Lancaster et al., 2014);
- “pI” – the predicted isoelectric point for a particular protein based on ExPASy (https://web.expasy.org/compute_pi/);
- “phasePlotRegion” – the region on the Das-Pappu diagram of states for a particular protein based on its sequence (Das and Pappu, 2013);
- “uversky hydropathy” – mean hydropathy as calculated from a skewed Kyte-Doolittle hydrophobicity scale (Kyte and Doolittle, 1982).
- “Ka” – non-synonymous evolutionary rates for the African killifish protein-coding genes
- “Ks” – synonymous evolutionary rates for the African killifish protein-coding genes for the African killifish protein-coding genes
- “Ka/Ks” – the ratio of non-synonymous over synonymous evolutionary rates.

To compute non-synonymous (Ka or dN) and synonymous (Ks or dS) evolutionary rates for the African killifish protein-coding genes, we first identified one-to-one orthologs among all the protein-coding genes in the African turquoise killifish, fugu, medaka, stickleback, tetraodon, and zebrafish using proteinortho (v5.15) (Lechler et al., 2017). We used the coding sequence corresponding to the longest isoform for alternative spliced genes. The coding sequences for the clusters that had a single orthologs for each of these species were aligned using prank (version v.140603) (Maiolo et al., 2018) and these codon-based alignments were filtered using GUIDANCE (version 2.0) (Sela et al., 2015). The Ka, Ks and Ka/Ks were computed using Yang and Nielsen algorithm implemented in Phylogenetic Analysis by Maximum Likelihood (PAML version 4.8) (Yang, 2007).

Because some of the sequence features did not conform to a normal distribution among detected proteins, we performed Monte Carlo sampling to determine the statistical significance of any enrichments or depletions. This analysis was performed on proteins that were aggregation-prone (with an aggregate abundance z-score >2, or roughly two standard deviations away from population mean, as a cutoff) and showed an age-associated increase in aggregation (paired t-test p-value <0.05 and OvY_logFC (log2 transformed fold change of AGG or PROP from old over young animals) z-score above 0.75). One-tailed statistical tests were performed for each tissue. Proteins identified in the tissue (either in the AGG or TL fraction) were sampled 10,000 times (with replacement) drawing exactly the same number of proteins as the query set to generate sample distributions. For the analysis on sequence properties for the aggregate fraction, the sampling of each protein was weighted by their relative abundance inferred from tissue lysate abundance in the young samples (sampling weight for a query protein was calculated as the ratio of total peptide spectrum matches of query protein to total peptide spectrum matches in tissue lysate sample) under the assumption that the null population can be modeled in a purely stochastic way dependent only on relative protein abundance. For analysis of sequence properties for the tissue lysate fraction and for aggregation propensity, samples of each protein were assigned equal weights (based on the assumption that there is no inherent abundance-dependent bias in sequence properties of most proteins). If the protein count information in the tissue lysate was not available, by definition, there was zero chance of sampling in the Monte Carlo simulation.

To infer if there was significant enrichment for properties of interest (except for PLACC score and Disordered Fraction), the p-value for each query protein set (i.e. aggregation prone proteins or age-associated aggregates) was obtained by calculating the fraction of samples that had a sample mean greater or smaller than query protein set. For normalized PLAAC score (“NLLR”), we used a minimum cutoff of 0 to classify protein as harboring a putative prion-like domain (∼7.0% of the killifish proteome and ∼6.8% of the detected killifish proteome met this cutoff respectively). Likewise, we used a threshold of 0.3 for Disordered Fraction (based on DISOPRED3 disorder) to determine whether a query protein is a putative intrinsically disordered protein (∼35.1% of the entire killifish proteome and ∼27.1% of the detected killifish proteome exceeded this cutoff). The p-values for enrichment of putative prions and intrinsically disordered proteins were obtained by calculating the fraction of samples that had higher prion-like or disordered fractions than the bona fide query protein set.

In Figure 3B, only those enrichments with a p-values below 0.05 were visualized as a non-gray square. The colors in the heatmap indicate the z-score of average feature value from the query set compared to randomly sampled test population mean, reflecting the extent of the enrichment of the indicated property.

#### Exogenous Expression of Age-associated Aggregation-prone Proteins in *S. cerevisiae*

Proteins with statistically significant (paired t-test p-value < 0.05) increase in aggregation propensity and/or aggregate abundance in old samples were cloned with custom DNA oligonucleotides (Table S19C) from killifish cDNA prepared from total RNA pooled from liver, brain, and muscle tissues. Briefly, liver, brain, and muscle tissues from male African turquoise killifish *N. furzeri* were individually homogenized in RLT buffer (RNeasy Kit, # 74104 QIAGEN) using 0.5 mm Silica disruption beads (RPI-9834) and a tissue homogenizer (FastPrep-24, 116004500 - MP Biomedicals). Total RNA was isolated from the lysed tissues according to the RNeasy Kit protocol. cDNA was prepared from the total RNA from pooled liver, brain, and muscle tissues with high-capacity cDNA RT kit (Applied Biosystems, 4368814) using random primers, and according to the manufacturers’ protocol. cDNA was amplified using custom DNA oligonucleotides (from IDT, Table S19C) and Phusion DNA Polymerase (Cat# F530L, Thermo Fisher Scientific), then cloned into a gateway entry vector (pDONR221, Cat# 12536017, Invitrogen). The resulting constructs were sequence verified against the annotated killifish genome (Table S1). The sequence-verified killifish ORFs were then cloned into a yeast gateway vector to allow GAL-inducible expression of the ORF with a C-terminal EGFP tag in yeast (pAG416GAL-ccdB-EGFP in Lindquist Advanced Gateway Vector Collection (Alberti et al., 2007)). The yeast expression plasmid (low-copy, CEN) for each protein was transformed into the standard laboratory strain BY4741. The strains bearing the plasmids were inoculated overnight in defined medium containing 2% raffinose as a carbon source (0.77 g of CSM-URA, 6.7 g of yeast nitrogen base without amino acid, and 20 g raffinose in 1 L media), then washed, diluted and switched to the same medium containing 2% galactose as a carbon source (0.77 g of CSM-URA, 6.7 g of yeast nitrogen base without amino acid, and 20 g galactose in 1 L media) to induce protein expression. The overnight culture generally reached an OD_600_ of 0.9-1 and was diluted to an OD_600_ of ∼0.1 prior to induction for 6-8 h, during which the OD_600_ of the cultures reached mid-exponential phase (OD_600_ ∼0.4-0.6). Microscopy images were all taken during mid-exponential phase using a Leica inverted fluorescence microscope with a Hammamatsu Orca 4.0 camera. Exposure times were 100 ms in the DIC channel and 50-500 ms in the fluorescent channel depending upon the signal strength of each GFP fusion protein (GFP excitation: 450–490 nm; emission: 500– 550 nm; software: LASX DMI6000B; refraction index: 1.518; aperture: 1.4; exposure time: 50, 250, and 500ms).

#### Support Vector Machine Classifier for Protein Aggregation State in Yeast

The support vector machine classifier was implemented in Python 2.7.15 using the sklearn module to identify the combination of parameters that most robustly separate the punctate and diffuse proteins in the yeast over-expression assay. We included our mass spectrometry data as well as the computed biophysical properties as features (the same features visualized in heatmaps in Figure 3B) to maximize the possible search space. Because the total number of experimentally tested proteins are small (47) compared to the number of possible features, we built a simple classifier that relies on fewer features to avoid overfitting. After transforming the features using standard score calculator (StandardScaler function, equivalent to z-score calculation), we iteratively tested two pairs of features at a time using a linear kernel function while fixing the regularization parameter C to 1. We performed cross-validation by splitting the dataset (20% of the data was reserved as test set), fitting a model, and computing the accuracy score 50 consecutive times. The score for each iteration for each pair is provided in Table S10 as well as the average score and the standard deviation. The OvY_prop_logFC (log2-transformed fold change of aggregation propensity in old over young animals) and charge patterning metric delta scored the best after cross-validation. The charge patterning metric delta in particular emerged as a key feature among the best performing two-feature classifiers we tested. The regularization parameter C for the two-feature linear support vector machine classifier went through further finetuning (vary C from 0.01 to 10) and the best hyperplane (with the highest accuracy score) that separates the two classes (diffuse and punctate proteins) among all proteins is plotted in Figure 4D.

#### Quantification of Cell Cycle Stages to Infer Tissue Proliferation Index

Three adult (2.5 months old) male transgenic killifish (*Nothobranchius furzeri*, GRZ strain) were used, carrying the cell-cycle dual FUCCI reporter (Dolfi et al., 2019) that allows for simple visualization of cell cycle using FACS. Fish were sedated in 200mg/L of Tricain (Sigma-Aldrich, A5040), and then euthanized in 500mg/L of Tricain in system water. Dissections were carried out under a stereo binocular (Leica S9E) at room temperature. Tissues (brain, gut, heart, liver, muscle, skin, and testis) were dissected from each fish and kept on ice in full medium (L15, 1% penicillin-streptomycin, 50 µg/µl gentamicin, 15% FBS). When all dissections were complete, organs were washed once with L15, and media was replaced with replaced with digestion media (400 µl 0.25% trypsin) at 28°C for 2 hours. After 2h, additional mechanical dissociation was applied by pipetting up and down with a 1ml tip for 15 minutes, followed by addition of 800µl of full media to stop enzymatic digestion. Dissociated cells were passed through a 100µm cell strainer prior to FACS analysis. For FACS analysis, cells from each tissue separately were stained with DAPI (0.1 µ g/ml), incubated for 15 minutes, and analyzed for GFP, RFP, and DAPI intensity using a CellStream™ analyzer FACS (Merck Millipore). Data analysis was performed with the integrates CellStream™ Acquisition and Analysis Software.

#### Disease Association Analysis

The disease association analysis was limited to proteins that were known to be associated with human Mendelian diseases (based on Online Mendelian Inheritance in Man, OMIM database downloaded on March 26, 2019). We focused our analysis on proteins with significant age-associated increase in aggregate abundance or aggregation propensity. A select group of proteins was highlighted and the abundance of these proteins in TL and AGG of young and old samples were visualized.

## Figure Generation

### Principal Component Analysis (PCA)

Principal Component Analysis (PCA) was performed in Python (version 2.7.15) using sklearn.decomposition.PCA function on the standardized log2-transformed normalized abundance for each protein in tissue lysate (TL) or protein aggregate (AGG) fractions (use sklearn.preprocessing.StandardScaler function for standardization) across different conditions (young, old, and old *TERT^Δ8/Δ8^* mutants).

### Seven-way Venn Diagram on Overlap Among Identified Proteins

This 7-way venn diagram was generated in R (version 3.5.1) using venn function in R package venn (version 1.7) on proteins detected in all seven tissues in tissue lysate (TL) and aggregate (AGG) fraction.

### Triangular Heatmap on Shared and Tissue-specific Changes Across Tissues

Tissue pair-wise comparison on shared and tissue-specific proteins identified in each category (proteins identified in TL or AGG, proteins with significant age-associated increase in TL/AGG/PROP where TL = tissue lysate abundance, AGG = aggregate abundance, PROP = aggregation propensity) were visualized in a heatmap. The number in each square represents the total number of shared proteins identified in the two tissues specified by the row label and column label.

### Heatmaps on Differential Changes in Tissue Lysate, Aggregate, and Aggregation Propensity

For each comparison (e.g. old vs young, old *TERT^Δ8/Δ8^* vs young, and old *TERT^Δ8/Δ8^* vs old), we used all the proteins (in TL, AGG, or PROP) that were significantly upregulated or down-regulated (p-value < 0.05) in at least one tissue. Proteins were sorted into two categories: 1) tissue-specific (proteins that had a significant up- or down-regulation in TL, AGG, or PROP in only a single tissue) at the top, or 2) shared (proteins that had a significantly up- or down-regulation in TL, AGG, or PROP in at least two tissues) at the bottom. The log2-transformed fold change results for the significant terms were colored in the heatmap with custom color bar (Figure 2A and S2C).

### Functional Enrichment Analysis

We identified enriched functional terms corresponding to Gene Ontology (GO), Diseases Ontology (DO), KEGG, KEGG-Modules and two MSigDb collections (Cellular Component and Hallmark Pathways) using Gene Set Enrichment Analysis (GSEA) implemented in R package clusterProfiler (version 3.10.1) (Yu et al., 2012). A separate enrichment analysis was performed on tissue lysate (TL) protein abundance, aggregate (AGG) protein abundance, and aggregation propensity (PROP) for each comparison (e.g. old vs young, old *TERT^Δ8/Δ8^* vs young and old *TERT^Δ8/Δ8^* vs old) for each tissue, and then combined and plotted based on shared functional terms. The protein lists were ranked and sorted in descending order based on multiplication of log2-transformed fold change and –log10(p-value). Note that due to random seeding effect in GSEA, the exact p-value and rank of the enriched terms may differ modestly for each run. This random seeding did not affect the enrichment analyses qualitatively.

### Heatmap on Functional Enrichment Analysis

Based on the functional enrichment analysis, we selected a short list of KEGG terms and visualized them as seen in Figure 2D. The top 3 significantly (p-value < 0.05) enriched Kyoto Encyclopedia of Genes and Genome (KEGG) terms with the highest normalized enrichment scores (NES) in every tissue were shown for TL, AGG, and PROP. Tissues-specific terms were placed on top whereas shared terms were placed at the bottom. The full lists of enrichment terms are available in Table S6. The color was scaled according to the rank statistic as computed by -log10(p-value) * log2(fold change).

### Analysis of Subcellular Localization

The cellular localization of killifish proteins was assumed to be similar to their human homologs. The assignment of killifish protein cellular compartment is available in Table S7A. Human protein localization information was retrieved from the Gene Ontology Consortium curated GO terms (downloaded on May 9, 2019). The GO annotations (go.obo) were first parsed into a table (go_obo_table.csv) where the GO term ID, name, namespace, definition, and parent GO term information were retained. Next, the human GO table (goa_human.gaf), which contains human protein IDs in UniProtKB and their associated GO terms IDs from various databases including GO and Reactome, was merged with GO annotation table (go.obo) so each human protein and all its corresponding GO terms information including name, namespace, and definition are available in one table. Because we use human Ensembl ID as unique identifier to map killifish proteins with their human orthologs (Table S2D), we retrieved the one-to-one map of Ensembl IDs to Uniprot IDs for each human protein from BIOMART and incorporated this to the human GO table (goa_human_ensembl.csv). We then assigned the cellular compartment of killifish proteins based on that of their human homologs (results in killifish_human_go.csv, available in GitHub repository associated with this manuscript). Because the GO terms compile information from multiple databases, there some redundancy among them. Therefore, to streamline the analysis, we primarily used the “cellular component” entries from Uniprot and Reactome as the other databases were less comprehensive and corresponded well with these two where there was overlap. Furthermore, we manually curated the cellular localization terms by using the more general classification (73 unique terms). For example, endoplasmic reticulum membrane and endoplasmic reticulum quality control compartment were combined into ‘endoplasmic reticulum’. The exact inclusion term and their classifications are available in reactome_CM_cleanup.csv for entries from Reactome and in uniprot_CM_cleanup.csv for entries from Uniprot. The final putative cellular compartment assignment of killifish proteins is available in Table S7A. The intermediate output is available in the GitHub repository (https://github.com/ywrchen/killifish-aging-aggregates) for this paper.

We first calculated the fraction of the observed proteome that is present in each subcellular location. If a protein was localized to multiple compartments, one count was assigned to each of them. To generate a comprehensive map of the subcellular localization of proteins that experienced age-associated changes in the TL and AGG fractions, we computed the fraction of proteins that reside in different cellular compartment for every issue. Results from cellular compartments that showed large tissue-specific differences were visualized in the donut plot in Figure 2E and Figure S2G. The quantification and visualization of cellular localization was performed in Python 2.7.15. In these donut plots, the reported percentage value (%) in the center is the average fraction of proteins that reside in the query compartment across tissues. If none of the proteins reside in a compartment for a given tissue, this tissue was not counted towards the calculation of average fraction and was omitted in the donut plot. The width of each slice for a tissue reflects the magnitude of the fraction (i.e. a tissue where more proteins come from a query compartment yields a larger slice of the donut).

### Charge Distribution Analysis and Visualization

The protein sequence feature analysis is described in “Biophysical and Sequences Features of Age-associated Aggregation-Prone Proteins” section. The “NCPR” (net charge per residue based on neighboring 5 amino acids, NCPR_blobLen5 in CIDER output) and hydropathy per residue (“hydropathy_blobLen5” based on the neighboring 5 amino acids) were obtained from localCIDER. Net positive charge and net negative charge residues were differentially colored along the protein sequence. A residue was considered hydrophobic if its hydropathy score exceeded 0.5 and was assigned a yellow bar for visualization. The disorder score profile was obtained from DISOPRED 3.

## SUPPLEMENTAL INFORMATION

Supplemental Information includes seven figures and 19 tables.

## SUPPLEMENTAL FIGURE LEGENDS

**Figure S1.**
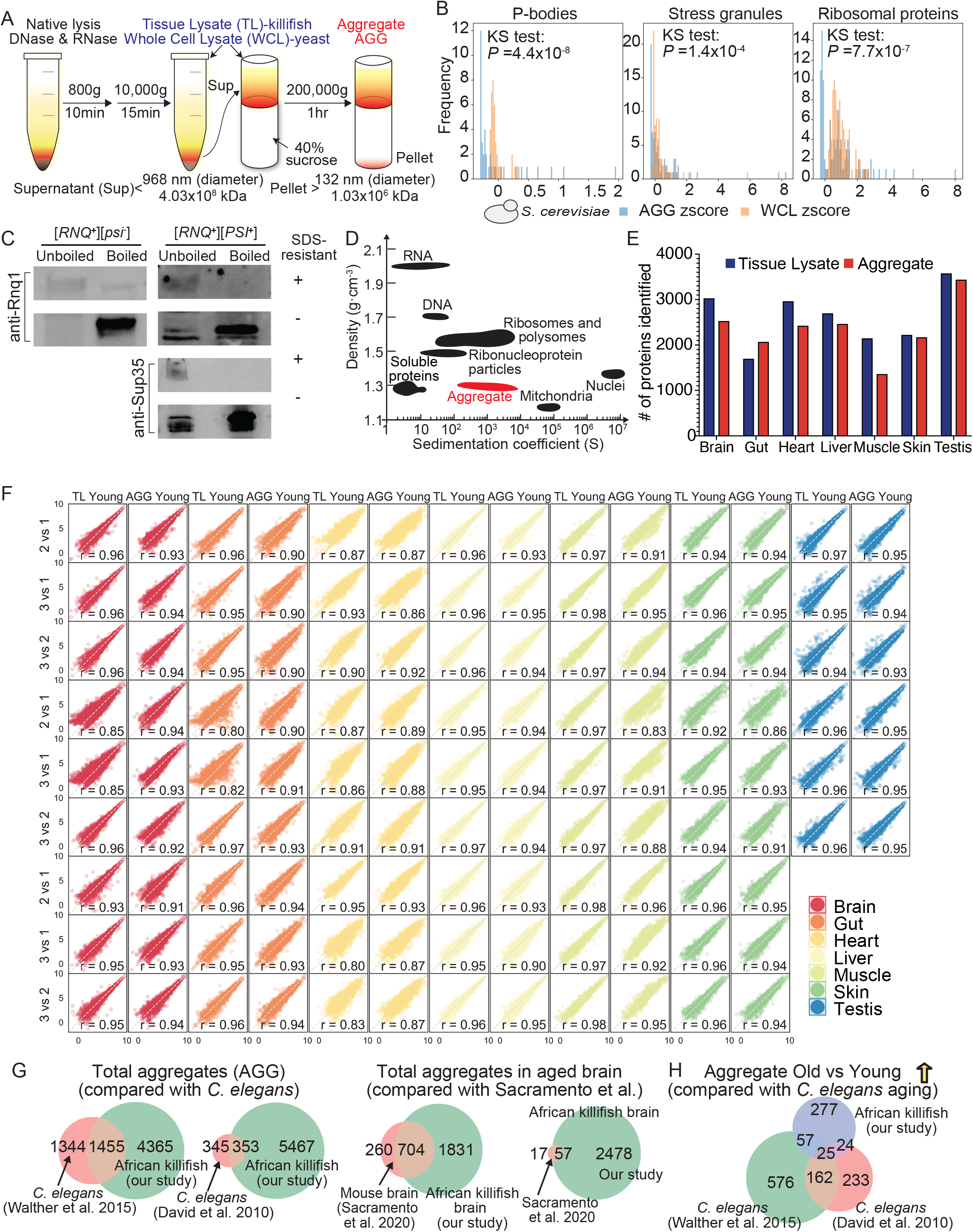
Related to Figure 1. Validation of high molecular weight aggregate isolation protocol in S. cerevisiae and quality control of tissue lysate and aggregate fractions in the African killifish. (A) Experimental workflow to extract total lysate – whole cell lysate (WCL) for yeast and tissue lysate (TL) for African killifish – and isolate high molecular weight protein aggregate fraction (AGG). S. *cerevisiae* pellet was lysed to extract whole cell lysate (WCL). The resulting yeast whole cell lysate was loaded onto a sucrose cushion and underwent ultracentrifugation to pellet high molecular weight fraction enriched with protein aggregates (AGG). Similarly, killifish tissues were homogenized to isolate tissue lysates (TL) and aggregates (AGG) and were subjected to ultracentrifugation with a sucrose cushion. The high molecular weight aggregate size was estimated to be 1.03 x 10^6^ – 4.03 x 10^8^ kDa in size and 132 – 968 nm in diameter (see STAR Methods for detailed theoretical calculations). (B) Distribution of the standardized protein abundance (represented as z-score in the x-axis) of processing bodies (left), stress granules (middle), and ribosomes (left) constituents identified in aggregate (AGG in blue, isolated from laboratory yeast strain BY4741 following the procedure described in A) and yeast total proteome (WCL in orange, calculated based on the unified proteome quantification of *S. cerevisiae* (Ho et al., 2018)). Two-sample two-sided Kolmogorov-Smirnov tests (KS-test) were performed to compare the two distributions for each category. (C) Western blot analysis of aggregates isolated from *S. cerevisiae* strains harboring known amyloid-forming prions ([*RNQ*^+^][*psi*^-^] (left) and [*PSI*^+^][*RNQ1*^+^] (right) 74D-695 strains). The isolated aggregate fractions (see A for isolation method) were resuspended in SDS sample buffer, and then divided in half. One half was boiled while the other was left un-boiled. Both were subsequently resolved on SDS-PAGE. SDS-resistant aggregates remained in (or near) the well while SDS-sensitive aggregates were denatured and migrated based on their respective molecular weights. Representative of 2 independent experiments. (D) Theoretically estimated densities (g. cm^-3^) and sedimentation coefficients (S) for the high molecular weight aggregates isolated using our protocol in comparison to those for nucleic acids (e.g., DNA, RNA), soluble proteins, protein complexes (e.g., ribosomes, polysomes, ribonucleoprotein particles such as spliceosome), organelles (mitochondria), and cellular compartment (nuclei). See STAR Methods for details of the calculation of density and sedimentation parameters for aggregates and comparison with theoretical sizes of other macromolecules or organelles in cells. (E) Mass-spectrometry coverage reported as the total number of proteins identified in tissue lysate (TL) and high molecular weight aggregate fractions (AGG) across all conditions (WT and *TERT^Δ8/Δ8^*) and age groups (young and old) for seven tissues. (F) Reproducibility of biological replicates within each age group for tissue lysate (TL) and aggregate fractions across tissues (AGG). Protein abundance (log2 transformed normalized peptide spectra counts) from respective biological replicates were plotted against each other. The Pearson’s correlation coefficient r is shown for each comparison and is also available in Table S2B. (G) Overlap between all aggregates identified in our dataset and previous studies that examined protein aggregation during aging in *C. elegans* (David et al., 2010; Walther et al., 2015) and aged African killifish and mouse brain (Kelmer Sacramento et al., 2020). Left: overlap in aggregates identified in our dataset (*C. elegans* homologs identified in young and old killifish samples from 7 tissues) and two *C. elegans* aging aggregate profiling studies (David et al., 2010; Walther et al., 2015)); Middle: overlap in aggregates identified in our dataset (mouse homologs and killifish proteins identified in old killifish samples from 7 tissues) and the aging vertebrate (mice and killifish) brain (Kelmer Sacramento et al., 2020). The identities of the shared total aggregate proteins are available in Table S3. (H) Overlap in aggregates with an age-associated increase in abundance (log2-transformed fold change of old over young sample log2FC > 0, two-sided Student’s t-test p-value < 0.05) among our killifish dataset and two *C. elegans* aggregate profiling studies (David et al., 2010; Walther et al., 2015). Age-associated aggregates from David et al. are proteins that consistently became 1.5-fold or more insoluble with age in all four datasets (Table S1 from David et al., 2010). Age-associated aggregates from Walther et al. are proteins identified from WT worms that showed increased aggregate abundance on day 17 compared to day 6 (Table S1D from Walther et al., 2015). The identities of the shared proteins and their age-associated changes in respective studies are available in Table S3.

**Figure S2.**
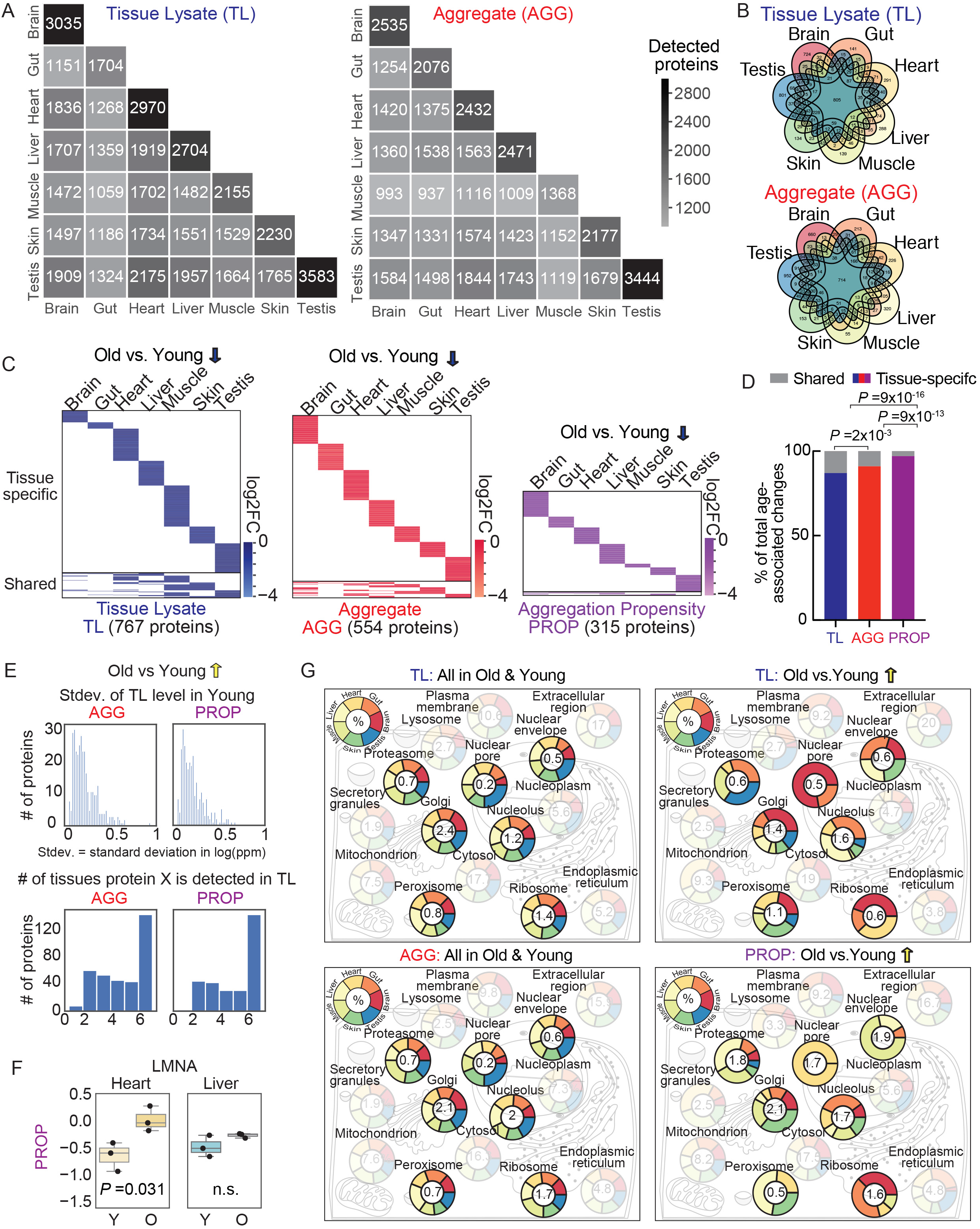
Related to Figure 2. Comparative analysis to probe the origin of tissue specificity in age-associated changes in tissue lysate, aggregates, and aggregation propensity. (A) Tissue pair-wise comparison on shared and tissue-specific proteins identified in TL or AGG. The number in each square represents the total number of shared proteins identified in the two tissues specified by the row label and column label. (B) Venn diagram of the identified proteins in tissue lysate and high molecular weight aggregate fractions across seven tissues. Note that the area size is not reflective of the actual number of proteins. (C) Heatmap on proteins that are significantly down-regulated (log2-transformed fold change in protein abundance log2FC < 0, two-sided Student’s t-test p-value < 0.05) with age (old over young) in tissue lysate (TL), aggregate (AGG), and aggregation propensity (PROP) across all tissues. Aggregation propensity is defined as the ratio of protein abundance in TL over AGG. The log2FC represents the average among 3 fish. Tissue-specific proteins (i.e., proteins with significant age-associated changes in only a single tissue) are at the top, and shared proteins (i.e., proteins with age-associated significant changes in at least two tissues) are at the bottom. The heatmaps are scaled based on the total number of proteins with such significant positive differential changes (shown in brackets for TL/AGG/PROP). The identities of proteins significantly down-regulated with age are in Table S5A-C. (D) Percentage of shared and tissue-specific changes among the entire dataset that showed a significant decrease (log2-transformed protein abundance fold change log2FC < 0, two-sided Student’s t-test p-value < 0.05) in tissue lysate (TL) protein abundance, high molecular weight aggregate fraction (AGG) protein abundance, or aggregation propensity (PROP) in old compared to young fish. Shared and tissue-specific proteins were defined as in Figure S2C. P-values are from a Chi-squared test of independence on tissue-specific versus shared proteins across all tissues for TL and AGG, TL and PROP, and AGG and PROP. Data are from Table S5D. (E) Quantification of variability in relative protein abundance (left: histogram of standard deviation among biological replicates in young killifish) across tissues and the extent of tissue-specific protein expression (right: histogram of the number of tissues a protein was detected in) among those with an age-associated increase in aggregate (top) or aggregation propensity (bottom). (F) Example of a protein with tissue-specific (only in heart) increase in aggregation propensity (PROP) during aging—driven by diverging changes in tissue lysate abundance (Figure 2C)—despite significant age-associated increase in aggregate (AGG) abundance in both tissues. P-values are from a Student’s t-test; n.s. indicates p>0.05. (G) Subcellular localization and complex association of proteins that are identified in tissue lysate (TL), aggregate fraction (AGG), as well as proteins with a significant age-associated increase in TL and aggregation propensity (PROP). The arc lengths in the doughnut plot for each tissue are proportional to the respective fractions of aggregates that increase with age and reside in the compartment across tissues. The average value of such fractions is reported in the center for each cellular compartment. Only tissues that contain proteins with a significant age-dependent increase in aggregates residing in the cellular compartments were visualized and counted towards the average calculation. Subcellular localization of killifish proteins was inferred from the homologous human proteins retrieved from UniProt localization database. Proteins that were annotated to exist in multiple compartments were double-counted. The fractions of upregulated aggregates residing in all compartments in each tissue are available in Table S7.

**Figure S3.**
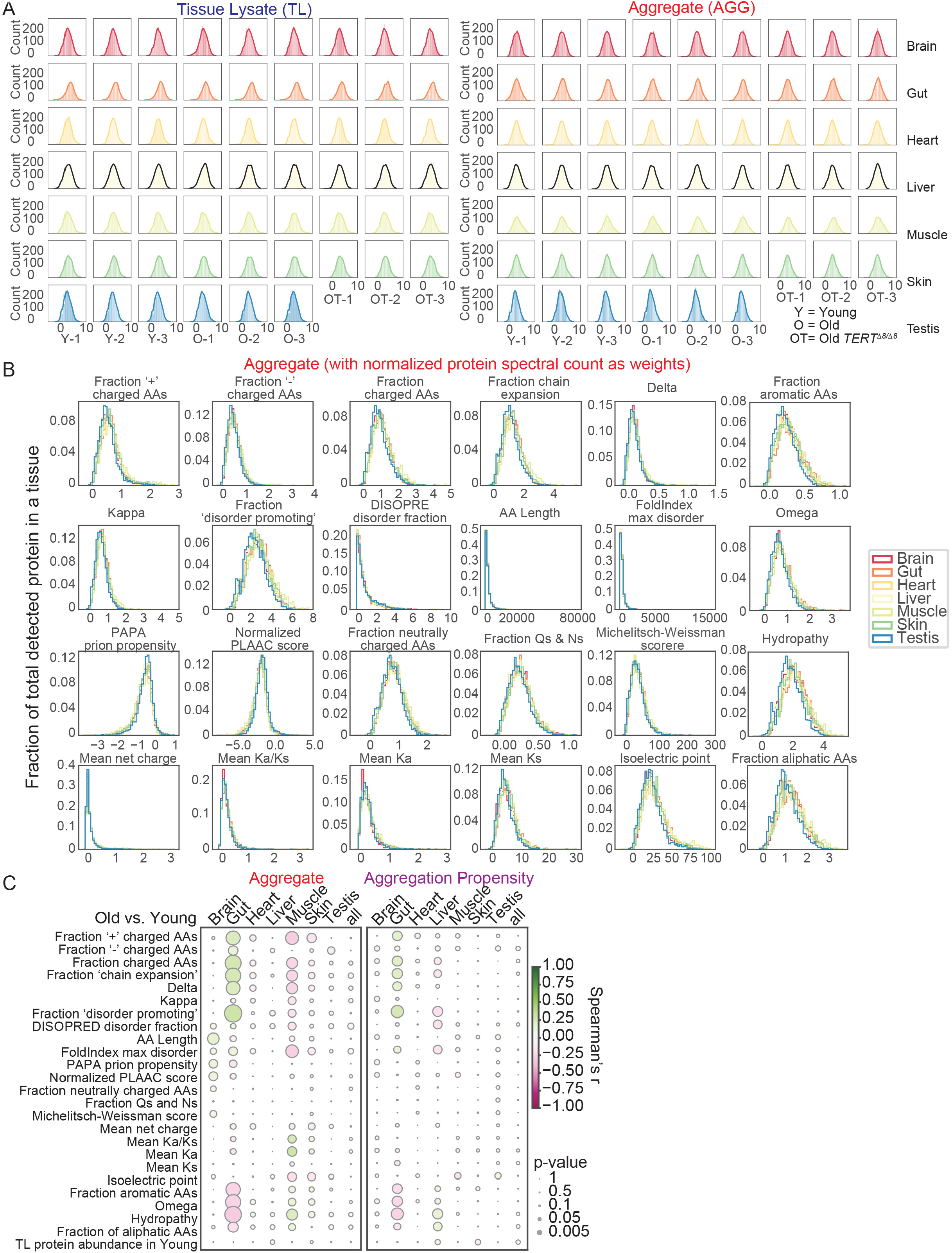
Related to Figure 3. Features of proteins detected in tissue lysate (TL) and aggregate (AGG) and regression analysis on age-associated changes in aggregate (AGG) and aggregation propensity (PROP) with these features. (A) Distribution of the log2-transformed protein abundances in tissue lysate (TL, left) and aggregate (AGG, right) in each killifish sample (young, old, and old *TERT^Δ8/Δ8^*). (B) Distribution of the biophysical features of all proteins detected in the high molecular weight aggregate fraction (AGG) of a tissue. Histograms of the products obtained from multiplying the value of protein sequence feature by the normalized protein abundance level (as weights) in tissue lysate (TL) are shown here. (C) The Spearman’s correlation coefficients from regression analysis of age-associated changes in aggregation (AGG) or aggregation propensity (PROP) against each query feature (all feature variables were continuous). The size of each circle is reflective of the p-value, and color is indicative of the magnitude of the correlation coefficient. The results are available in Table S8E-F.

**Figure S4.**
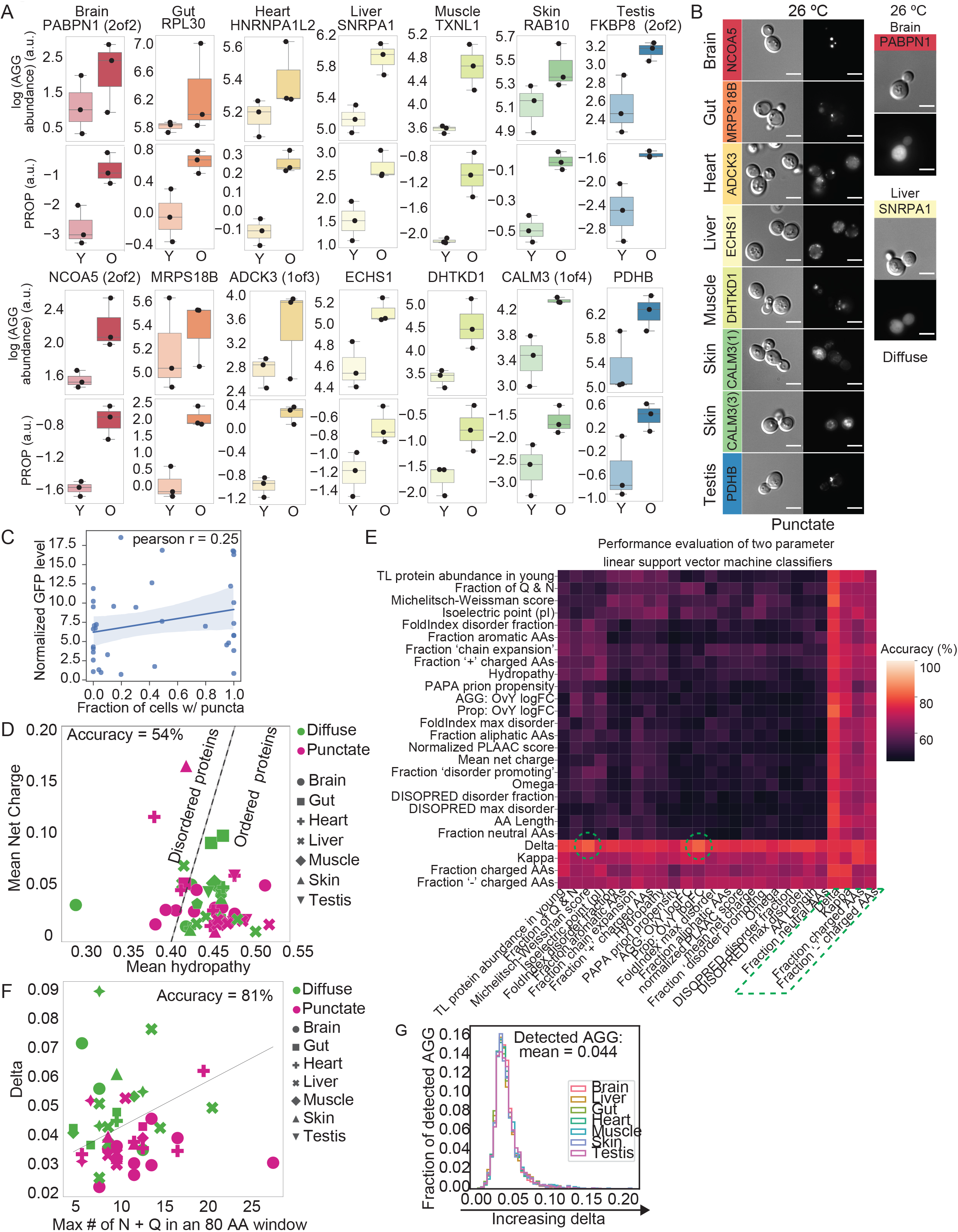
Related to Figure 4. Age-associated changes in aggregate and aggregation propensity of proteins tested in yeast overexpression assay and assessment of *in vivo* aggregation behavior predictor. (A) Box plot representation of the aggregate abundance and aggregation propensity levels in young and old animals on selected proteins that were tested in *S. cerevisiae* overexpression assay. The box shows the quartiles of the data while the whiskers extend to show the rest of the distribution on protein abundance (y-axis) in young (Y) or old (O) samples. Each dot represents the AGG abundance or PROP level from an individual fish. (B) Example microscopy images of the proteins of interest (with a significant age-associated increase in aggregate and/or aggregation propensity in old versus young animals) upon overexpression in *S. cerevisiae* as outlined in Figure 4C except that the yeast was cultured at 26 °C. Two diffuse proteins were tested and shown on the right. One punctate protein example was chosen and shown for each tissue on the left. The images chosen were representative of two independent experiments. (C) Scatter plot on GFP expression level and the fraction of GFP positive cells with puncta tested in *S. cerevisiae* overexpression assay. Each dot represents result from a strain overexpressing a killifish protein of interest. The GFP expression level is a normalized value, approximated by subtracting background GFP intensity from inside GFP positive cells followed by division of background intensity, based on GFP intensity measured by fluorescent microscopy. (D) Uversky plot (mean net charge versus mean hydropathy) on age-associated aggregates tested in the *in vivo* aggregation assay. The dashed line represents the boundary defined by and <H> = (<R> + 1.151)/2.785, where <H> is the mean hydropathy and <R> is the mean net charge of a protein sequence. Proteins to the left of the dotted line are predicted to be ordered, whereas those to the right are predicted to be unstructured and disordered. (E) Performance evaluation of two-parameter linear support vector machine classifiers in predicting proteins aggregation behavior *in vivo* based on our proteomics study and protein biophysical properties. All the trained models and their performance are available in Table S10. See STAR Methods for details. (F) Support vector machine classifier based on the Michelitsch-Weissman score (maximum number of glutamine (Q) and asparagine (N) residues within an 80 amino acid windows of a protein) and charge patterning metric “delta” for proteins tested in yeast autonomous aggregation assay. *In vivo* aggregation status was distinguished by color. Tissues in which the protein exhibited increased aggregation propensity changes can be distinguished by the different marker shapes. The dotted line separates those that showed diffuse eGFP morphology upon protein overexpression in *S. cerevisiae*. (G) Histogram of charge patterning metric “delta” of aggregate detected in all seven tissues.

**Figure S5.**
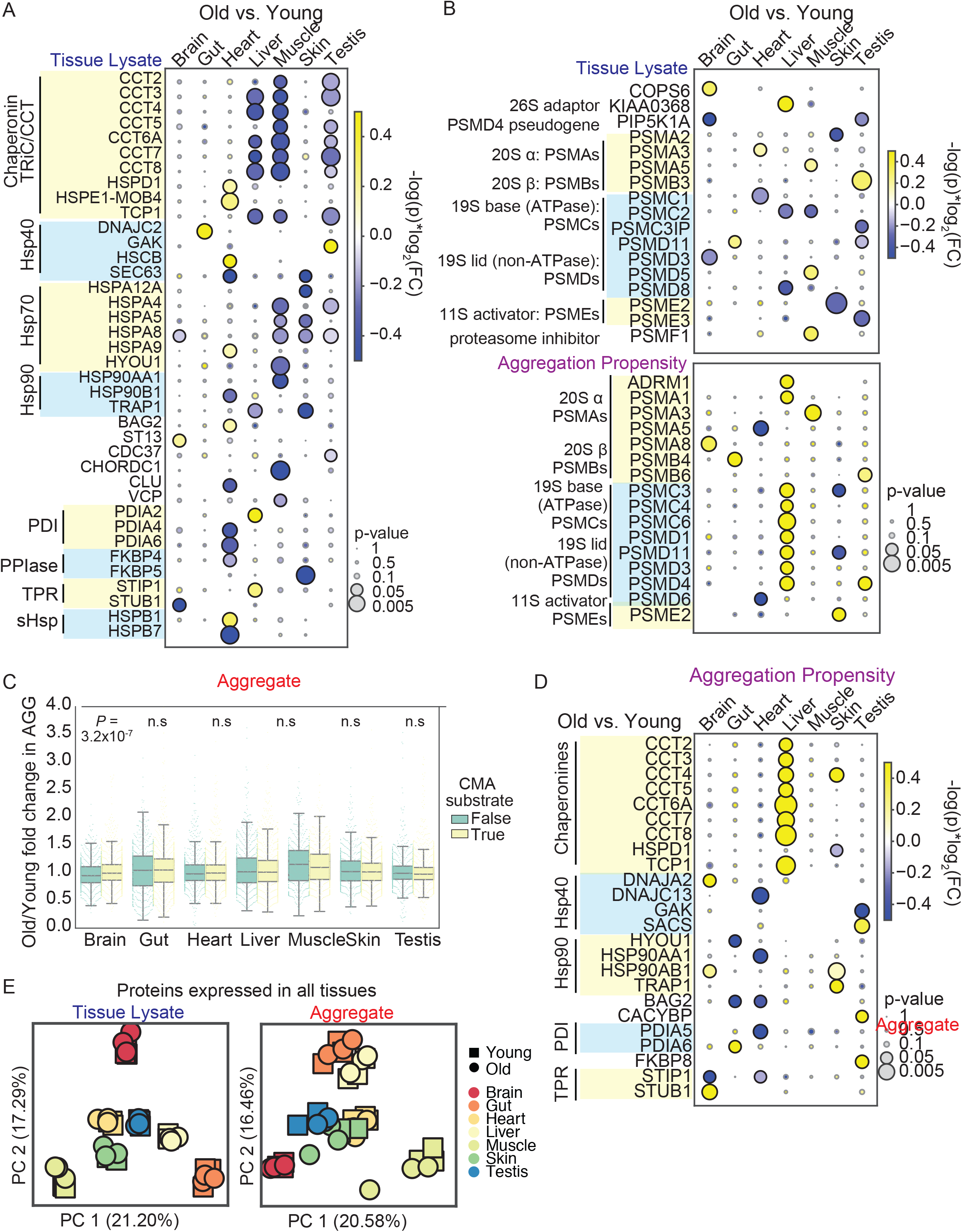
Related to Figure 5. Additional examples of tissue-specific age-associated changes in protein quality control machinery. (A) Chaperone abundance changes in tissue lysate between old and young animals. The circle color is indicative of the ranked statistics (-log10-transformed p-value multiplied by log2-transformed fold change of old over young protein abundance), and the circle size is indicative of the -log10-transformed p-value from Student’s t-test. The chaperones were grouped by their classes. Results are available in Table S11. (B) Tissue lysate (TL) and aggregation propensity (PROP) changes of proteasome component with age. The circle color is indicative of the ranked statistics (-log10-transformed Student’s t-test p-value multiplied by log2-transformed fold change of old over young protein abundance), and the circle size is indicative of the -log10-transformed Student’s t-test p-value. Results are available in Table S11. (C) Aggregate abundance changes of CMA-selective clients versus non-client in aging killifish. CMA-clients are proteins detected in killifish brain with peptide motif that meet the following criteria (Kaushik and Cuervo, 2018): include glutamine on one of the sides and contains one or two of the positive residues K and R, one or two of the hydrophobic residues F, L, I or V and one of the negatively charged E or D residues. Fold change in aggregate abundance of clients and non-clients in old versus young killifish brain were separately plotted (each dot represents a single detected protein). A boxplot is overlayed to show the quartiles of the dataset while the whisker extends to show the rest of the distribution. Independent two-sided t-tests were performed between clients and non-client for each tissue, and the p-values were reported. (D) Aggregation propensity changes in chaperones with age. The circle color is indicative of the ranked statistics (-log10-transformed Student’s t-test p-value multiplied by log2-transformed fold change of old over young), and the circle size is indicative of the -log10-transformed Student’s t-test p-value. Results are available in Table S11. (E) Principal component analysis on protein abundance of proteins detected in all seven tissues in tissue lysate and aggregate fractions. Each symbol represents a sample from an individual fish. The shape of each marker indicates young (squares) and old (circles) fish, and the color indicates the tissue origin.

**Figure S6.**
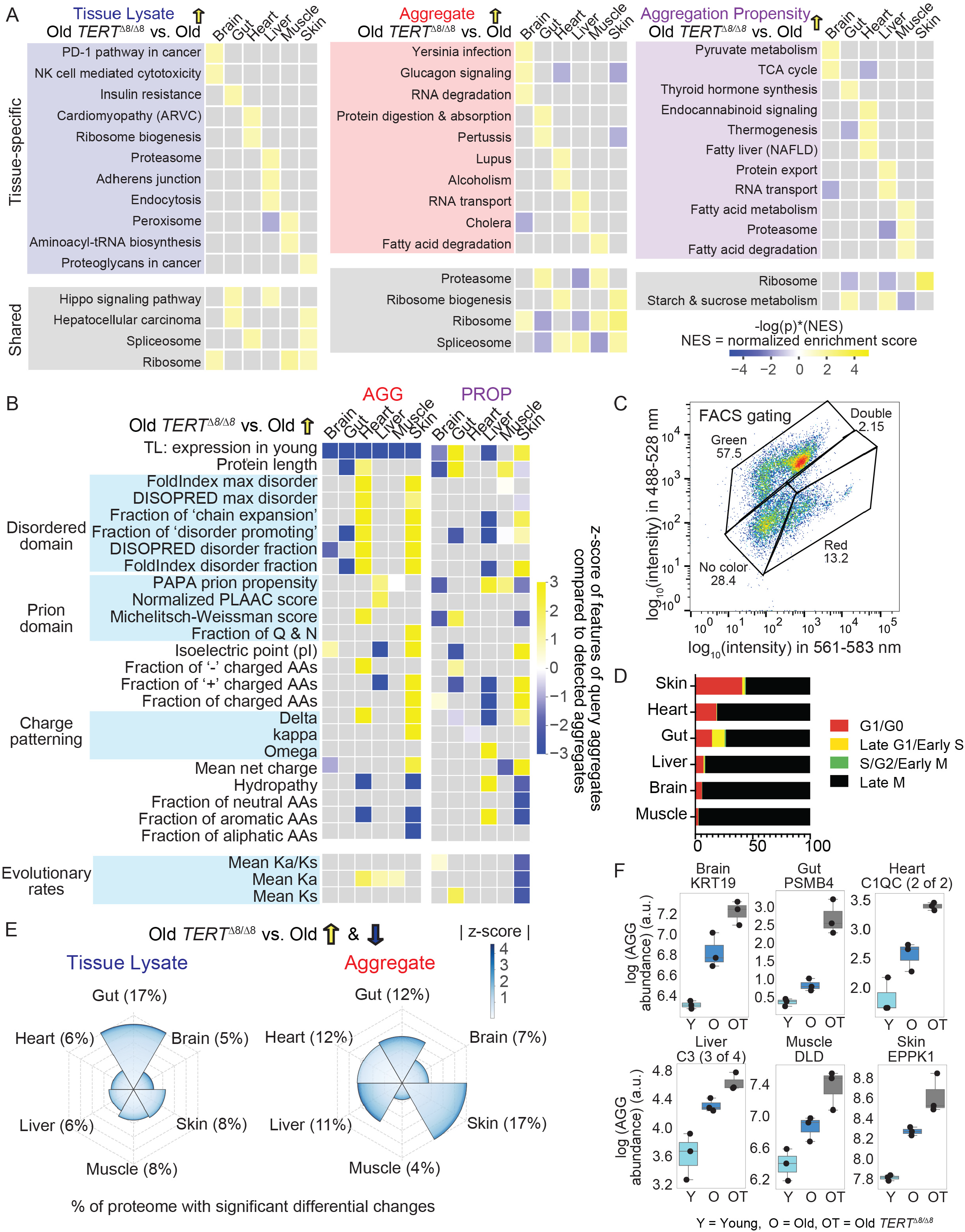
Related to Figure 6. Disease-associated changes in old *TERT*^Δ8/Δ8^ mutant compared to age-matched wild-type animals. (A) Heatmap of significant functional and pathway enrichments identified among upregulated proteins in old *TERT^Δ8/Δ8^* versus age-matched wild-type killifish using Gene Set Enrichment Analysis (GSEA). The proteins were ranked and sorted in descending order based on multiplication of log2-transformed fold change and –log10(p-value) (yellow upward arrow). Due to space constraints, only the top 3 (ranked by the highest normalized enrichment scores (NES)) significantly (p-value < 0.05) enriched Kyoto Encyclopedia of Genes and Genome (KEGG) terms in every tissue are shown for TL, AGG, and PROP. Tissues-specific terms are placed on top, whereas shared terms are placed at the bottom. The full lists of enrichment terms are available in Table S14. The color is scaled according to the rank statistic computed as the product of multiplying - log10(p-value) by NES. (B) Biophysical features of proteins with significant (Student t-test p-value < 0.05) increase in aggregate (AGG) and aggregation propensity (PROP) in old *TERT^Δ8/Δ8^* versus age-matched wild-type killifish. Terms were visualized only when the average feature value from the query set of proteins were significantly (one-tail test at 0.05 cutoff) different from the Monte Carlo simulation derived population distribution, whereas the nonsignificant ones were in gray. The color in the heatmap reflected the z-score of average feature value from the query set compared to Monte Carlo simulation derived test population means. See STAR Methods for details on the implementation and analysis of the Monte Carlo simulation. (C) Example of FACS gating based on GFP and RFP intensity applied to three young (12-week-old wild-type killifish stably integrated with FUCCI reporter) testis samples. Quantification of different cell cycle stages in each tissue is available in Table S14. (D) Quantification of different cell cycle stages assessed by FUCCI reporter cell line described in Figure 6C. The results are average from 3 biological replicates. All results are available in Table S14. (E) Percentage of proteins with differential changes (include both up-regulation and down-regulation) in tissue lysate (TL) and aggregate (AGG) between age-matched old *TERT*^Δ8/Δ8^ and wild-type animals. The z-scores of changes in TL or AGG between age-matched old *TERT^Δ8/Δ8^* and wild-type were calculated for each protein (i.e., TvO_prop_logFC was the log2-transformed fold change of old *TERT^Δ8/Δ8^* divided by age-matched wild-type) at each tissue level. Those that were significant (p-value <0.05) were counted, and the percentage of proteome they represent is reflected in the radius of the sector for each tissue. The heatmap color is reflective of the ranked z-score of these significantly differentially regulated proteins with a higher degree of remodeling denoted with darker shades of blue. (F) Examples of age-associated aggregates with further enhanced aggregate burden upon telomerase deficiency (proteins with increased aggregate burden between old and young animals, as well as age-matched old *TERT*^Δ8/Δ8^ and old wild-type). The box shows the quartiles of the data while the whiskers extend to show the rest of the distribution on protein abundance (y-axis) in old (O) or old *TERT^Δ8/Δ8^* (OT) samples. Each dot represents the TL or AGG abundance from an individual fish.

**Figure S7.**
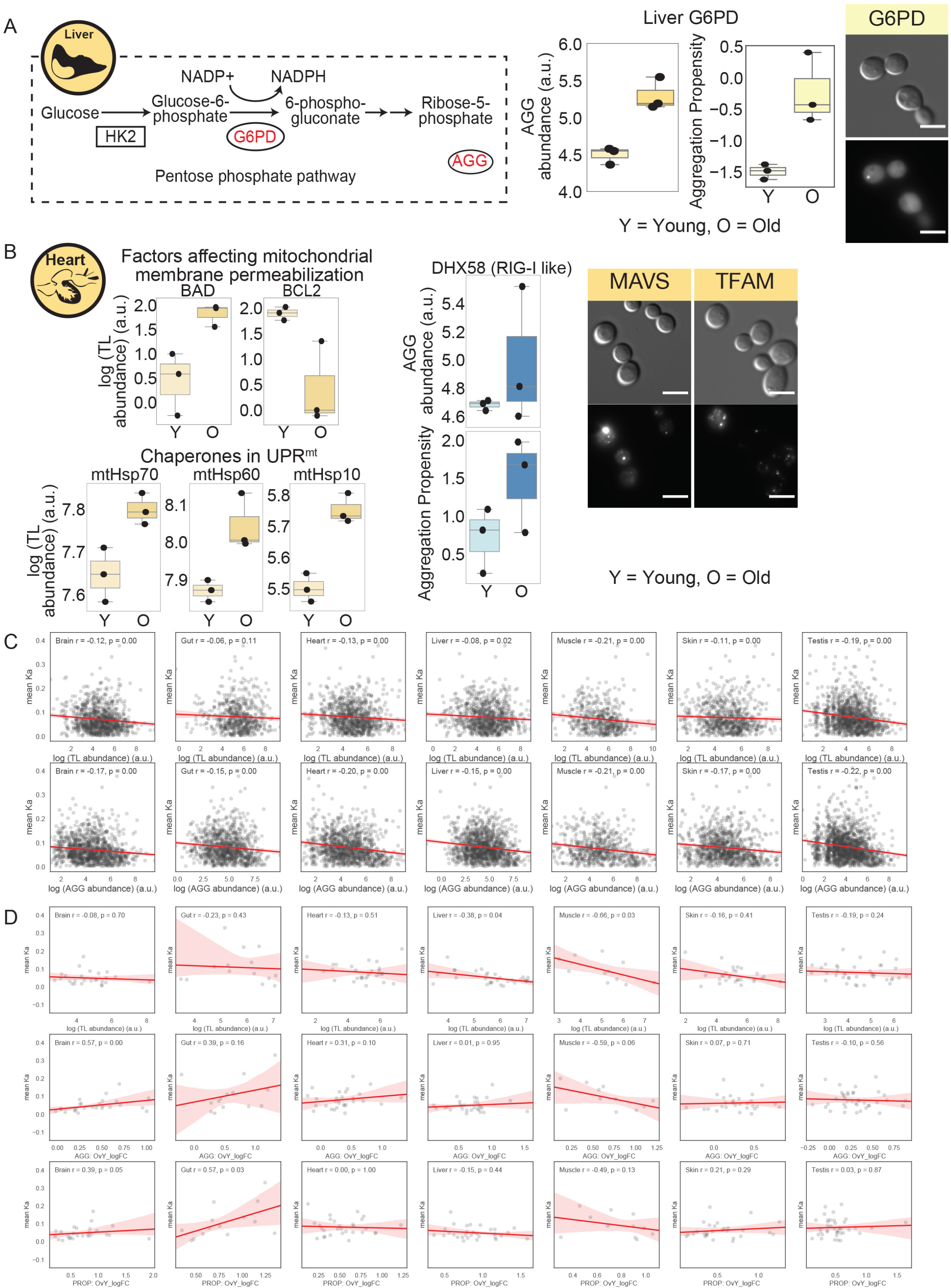
Related to Figure 7. Other age-associated aggregates implicated in diseases including those not previously linked to protein misfolding. (A) Network diagram of G6PD catalyzed reactions and box plot representation of its aggregate and aggregation propensity level in young and old killifish liver. G6PD is not known to aggregate previously. The box shows the quartiles of the data while the whiskers extend to show the rest of the distribution on protein abundance (y-axis) in old (O) or old *TERT^Δ8/Δ8^* (OT) samples. Each dot represents the AGG abundance or PROP value from an individual fish. (B) Box plot on tissue lysate changes in a few mitochondrially localized proteins and box plot of aggregate and aggregation propensity changes of the putative killifish RIG-I in young and old killifish heart. The box shows the quartiles of the data while the whiskers extend to show the rest of the distribution on protein abundance (y-axis) in young (Y) and old (O) samples. Each dot represents the TL abundance or AGG abundance or PROP value from an individual fish. (C) Scatter plot between non-synonymous mutation rate (Ka, y-axis) and TL abundance (top row, x-axis) or AGG abundance (bottom row, x-axis) level in young killifish tissues. (D) Scatter plot between non-synonymous mutation rate (Ka, y-axis) and TL abundance (top row, x-axis), fold change in aggregate abundance between old and young fish (middle, x-axis), and fold change in aggregation propensity between old and young fish (bottom row, x-axis) level among proteins with significant up-regulation (p-value <0.5, fold change > 0) in aggregation propensity during aging.

## Notes

### Competing Interest Statement

The authors have declared no competing interest.

